# A temporally resolved, multiplex molecular recorder based on sequential genome editing

**DOI:** 10.1101/2021.11.05.467388

**Authors:** Junhong Choi, Wei Chen, Anna Minkina, Florence M. Chardon, Chase C. Suiter, Samuel G. Regalado, Silvia Domcke, Nobuhiko Hamazaki, Choli Lee, Beth Martin, Riza M. Daza, Jay Shendure

## Abstract

DNA is naturally well-suited to serve as a digital medium for *in vivo* molecular recording. However, DNA-based memory devices described to date are constrained in terms of the number of distinct signals that can be concurrently recorded and/or by a failure to capture the precise <u>order</u> of recorded events^1^. Here we describe DNA Ticker Tape, a general system for *in vivo* molecular recording that largely overcomes these limitations. Blank DNA Ticker Tape consists of a tandem array of partial CRISPR-Cas9 target sites, with all but the first site truncated at their 5’ ends, and therefore inactive. Signals of interest are coupled to the expression of specific prime editing guide RNAs^2^. Editing events are insertional, and record the identity of the guide RNA mediating the insertion while also shifting the position of the “write head” by one unit along the tandem array, *i.e.* sequential genome editing. In this proof-of-concept of DNA Ticker Tape, we demonstrate the recording and decoding of complex event histories or short text messages; evaluate the performance of dozens of orthogonal tapes; and construct “long tape” potentially capable of recording the order of as many as 20 serial events. Finally, we demonstrate how DNA Ticker Tape simplifies the decoding of cell lineage histories.

## Introduction

How do we learn the order of molecular events in living systems? A first approach is direct observation, *e.g.* live cell fluorescence microscopy to quantify the interactions in real time. A second approach is time-series experiments, *e.g.* destructively sampling and transcriptionally profiling a system at different timepoints. A third approach is epistatic analysis, *e.g.* ordering the actions of genes by comparing the phenotypes of single and double mutants. Although these and other approaches have important strengths, they are also limited in key ways. For example, live imaging is largely restricted to *in vitro* models. For time series experiments, resolution and accuracy are constrained by the frequency of sampling and the reproducibility of the biological process under investigation. Epistatic analysis is confounded by pleiotropy, particularly in multicellular organisms.

Another approach to ordering molecular events, theoretically promising but methodologically underdeveloped relative to the aforementioned alternatives, is a DNA memory device^3^, which we define here as an engineered system for digitally recording molecular events through permanent changes to a cell’s genome that can be read out in *post hoc* fashion. To date, several proof-of-concept DNA memory devices have been described that leverage diverse approaches for the “write” operation, including site-specific recombinases (SSRs)^4,5^, CRISPR-Cas9 genome editing^6–9^, CRISPR integrases^10,11^, terminal deoxynucleotidyl transferases^12^, base-pair misincorporation^13^, base editing^14^, and others^1^.

The nature of the write operation in such molecular recorders shapes their performance in terms of channel capacity for encoding and decoding signals, temporal resolution, interpretability and portability^1^. For example, SSRs record molecular signals with high efficiency, but the number of distinct signals that can be concurrently recorded is limited by the number of available SSRs. Molecular recorders relying on CRISPR-Cas9 can potentially overcome this limitation, *e.g.* if each signal of interest were coupled to the expression of a different guide RNA (gRNA), but in that case each signal would also require its own target(s). Furthermore, the CRISPR-Cas9 molecular recorders described to date rely on double-stranded breaks (DSBs) and nonhomologous end-joining (NHEJ) to write events to DNA^1^. Frequent DSBs are toxic and often result in the deletion of any consecutively located target sites, the molecular equivalent of accidental data deletion.

A further handicap of nearly all DNA memory devices described to date is that events are stochastically recorded to unordered target sites, thereby obscuring the order in which they occurred. CRISPR integrase systems, which rely on the signal-induced, unidirectional incorporation of DNA spacers or transcript-derived tags to an expanding CRISPR array, overcome this limitation^10,11,15–17^. However, at least to date, their reliance on accessory integration host factors has restricted such recorders to prokaryotic systems. Another recently described approach to enable directional writing of information to DNA combines self-targeting CRISPR gRNAs with the expression of terminal deoxynucleotidyl transferase (TdT), whose presence shifts the most likely outcome of NHEJ from short deletions to short insertions^18^. While this approach unidirectionally inserts sequences in a signal-responsive manner, it continues to rely on toxic DSBs, and because each gRNA/target yields a homogenous signal (*i.e.* TdT-mediated insertions of variable length), it is not clear how it could be used to explicitly record the precise order of a very large number of distinct signals.

Here we describe a DNA-based memory device that is: (1) highly multiplexable, *i.e.* compatible with the concurrent recording of at least thousands of distinct signals; (2) sequential and unidirectional in recording events to DNA, and therefore able to explicitly capture the precise temporal order of recorded events; (3) active in mammalian cells. This system, which we call DNA Ticker Tape, begins with a tandem array of partial CRISPR-Cas9 target sites, all but the first of which are truncated at their 5’ ends, and therefore inactive (**Figure 1a**). Each signal of interest is coupled to the expression or activity of a prime editing guide RNA (pegRNA)^2^. Alternatively, for some applications (*e.g.* lineage tracing), pegRNA expression could be constitutive. These pegRNAs, together with the prime editing enzyme^2^, are designed to mediate the insertion of a k-mer within the sole active site of the tandem array, which is initially its 5’-most target site. In the simplest implementation, all pegRNAs target the same 20-bp spacer, but each encodes a distinct k-mer insertion. Specifically, the 5’ portion of the k-mer insertion is variable and encodes the identity of the pegRNA, while the 3’ portion is constant, and activates the subsequent target site in the tandem array by restoring its 5’ end. Thus, each successive edit records the identity of the pegRNA that mediated the edit, while also shifting the position of the active target site by one unit along the array. At any moment, an intact spacer and PAM are present at only one location along the array, analogous to the “write head” of a disk drive.

**Figure 1.**
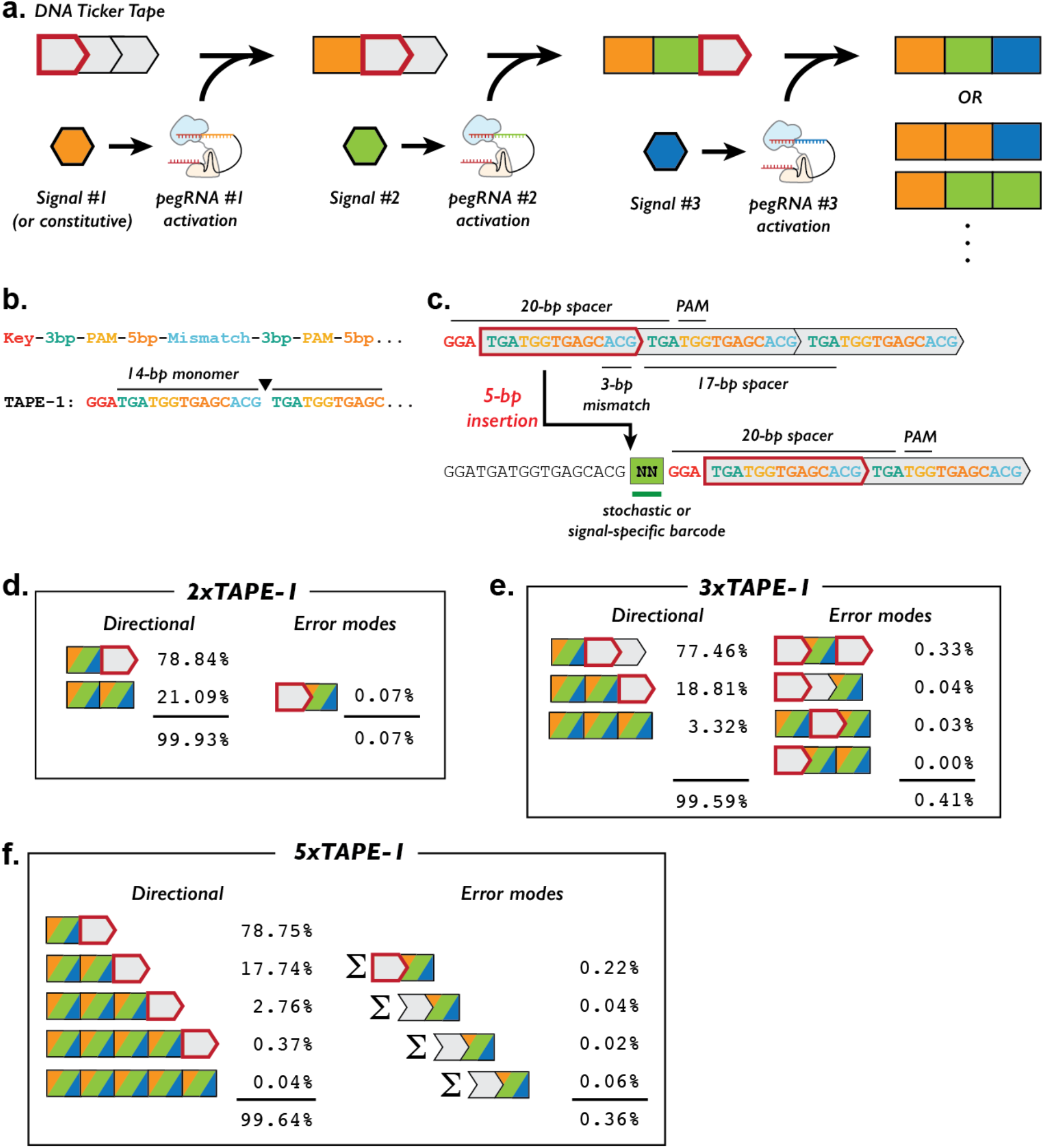
Sequential genome editing with DNA Ticker Tape. **a.** Schematic of DNA Ticker Tape. Individual pegRNAs are either signal-driven or constitutively expressed, together with the PE2 enzyme. The tape consists of a tandem array of CRISPR-Cas9 target sites (grey boxes), all but the first of which are truncated at their 5’ ends, and therefore inactive. Editing of the sole active site (red outline) inserts a pegRNA-specific barcode as well as a “key” sequence that activates the next monomer along the array. Because genome editing is sequential in this scheme, the temporal order of recorded events can simply be read out by their physical order along the array. **b.** TAPE-1 is a modified version of the HEK3 prime editing target sequence and consists of a key (GGA), followed by tandem repeats of a 14-bp monomer. The 14-bp monomer includes a PAM at positions 4-6 as well as mismatches to the key at positions 12-14. **c.** Schematic of editing event at the “write head” of TAPE-1. The 5-bp insertion includes a 2-bp pegRNA-specific barcode as well as a 3-bp key that activates the next monomer. **d-f.** Specificity of genome editing on versions of TAPE-1 with two (**d**), three (**e**), or five (**f**) monomers. Cells bearing stably integrated TAPE-1 target arrays were transfected with a pool of plasmids expressing pegRNAs and PE2. The TAPE-1 region was then PCR-amplified and sequenced. For each class of editing pattern, we report its proportion out of total reads with 1+ edits, which were 5.3%, 7.2%, and 6.6% of all reads for 2xTAPE-1, 3xTAPE-1, and 5xTAPE-1, respectively. Each class of outcomes is inclusive of all possible NNGGA insertions, and collectively the classes shown include (2^n^ - 1) possible outcomes, where *n* is the number of monomers. As predicted, we observe that editing of any given target site is highly dependent on the preceding sites in the array having already been edited (*e.g.* with 5xTAPE-1, of the reads in which Site-3 was edited, 98.8% were also edited at both Site-1 and Site-2).

## Proof-of-concept of DNA Ticker Tape

To test this idea, we designed a DNA Ticker Tape (“TAPE-1”) by modifying a spacer sequence previously shown to be highly amenable to prime editing by the PE2 enzyme (HEK293 target 3 or *HEK3*)^2^. In TAPE-1, a 3-bp “key” (GGA) is followed by one tandem array of a 14-bp monomer (TGATGGTGAGCACG) that includes the PAM sequence (TGG) at positions 4-6. At the 5’-most end of the TAPE-1 array, the key sequence, the first 14-bp monomer, and the first 6 bases of the subsequent 14-bp monomer, collectively comprise an intact 20-bp spacer and PAM (**Figure 1b**). We further designed a set of 16 pegRNAs to target TAPE-1, with each pegRNA programming a distinct 5-bp insertion. The first 2-bp of the insertion was unique to each of the 16 pegRNAs. The remaining 3-bp of the insertion corresponded to the key (GGA). We reasoned that when a pegRNA/PE2-mediated insertion occurred at the active TAPE-1 site, it would: (1) record the identity of the pegRNA -- via the 2-bp portion of the insertion; (2) inactivate the current active site -- by disrupting its sequence; and (3) activate the next monomer along the array -- as the newly inserted GGA key, together with the subsequent 20-bp, creates an intact 20-bp spacer and PAM (**Figure 1c**). In the next iteration of genome editing, a pegRNA-mediated insertion to the second monomer would be recorded while also moving the “write head” to the third monomer, and then to the fourth, the fifth, and so on.

We synthesized and cloned TAPE-1 arrays with varying numbers of monomer units (2xTAPE-1, 3xTAPE-1, 5xTAPE-1), and stably integrated these arrays into the genome of HEK293T cells via the piggyBAC system. We transiently transfected the resulting cells with a pool of plasmids designed to express PE2 and sixteen pegRNAs, each programmed to insert an NNGGA barcode to TAPE-1, and harvested them after four days. The TAPE-1 region was PCR-amplified from genomic DNA and sequenced.

For each TAPE-1 array, we categorized sequencing reads into those in which: (1) no editing occurred; (2) the observed pattern was consistent with sequential, directional editing; or (3) the observed pattern was inconsistent with sequential, directional editing (**Figure 1d-f**; **Supplementary Table 1**). Overall editing rates were modest, as only 5.3%, 7.2%, and 6.6% of all reads for 2xTAPE-1, 3xTAPE-1, and 5xTAPE-1, respectively, exhibited any editing. However, within the set of reads showing edits, the data were overwhelmingly consistent with sequential, directional editing. For example, with 2xTAPE-1, the second monomer was edited in 21.1% of reads in which the first monomer was also edited (**Figure 1d**). In contrast, the second monomer was only edited in 0.004% of reads in which the first monomer was <u>not</u> edited. This observation strongly suggests that edits of the second monomer were dependent on an edit of the first monomer having already occurred. It further confirms that the 3-bp mismatch at the PAM-distal end of “inactive” spacers of the TAPE-1 design is sufficient to inhibit prime editing. Data obtained from 3xTAPE-1 and 5xTAPE1 were also consistent with sequential genome editing. For example, 98.5% (3xTAPE-1) and 99.0% (5xTAPE-1) of reads that were edited at the second monomer were also edited at the first monomer, while 97.6% (3xTAPE-1) and 98.8% (5xTAPE-1) of reads that were edited at the third monomer were also edited at the first and second monomers (**Figure 1e-f**). These results were consistent across three transfection replicates (**Supplementary Table 1**).

An interesting phenomenon is that while the observed editing rate of the first TAPE-1 monomer was around 5-7%, the editing rates of the second or third TAPE-1 monomers, given that the preceding monomers were already edited, were around 15-25%. A simple explanation for the roughly three-fold greater “elongation” than “initiation” of editing is that some arrays are less amenable to prime editing than others by virtue of their site of integration, resulting in an excess of fully unedited arrays. Factors such as heterogeneous susceptibility of cells to transfection and the influence of cell cycle phase on editing efficiency might also contribute to this “pseudo-processivity”.

We next analyzed the distribution of the 16 NNGGA barcode insertions, focusing on 5xTAPE-1 (**Supplementary Figure 1**). Their frequencies correlated well amongst three replicates as well as between the first and second target sites (Pearson’s *r* = 0.97-0.99; **Supplementary Figure 1a-b**). The observed variation was partly explained by the relative abundances of the individual pegRNAs in the plasmid pool (Pearson’s *r* = 0.87; **Supplementary Figure 1c**). To explore whether the sequence of the insertion itself influences editing efficiency, we repeated the experiment but with an equimolar pool of 16 pegRNA-expressing plasmids that had been individually cloned and purified (rather than cloned as a pool). For each of the NNGGA insertions in each experiment, we calculated “edit scores” as log-scaled insertion frequencies normalized by abundances of pegRNAs in the corresponding plasmid pools. These edit scores were well correlated between 5xTAPE-1 edited by the 16 pegRNA plasmids pooled pre- vs. post-cloning (Spearman’s *p* = 0.97; **Supplementary Figure 1d**), consistent with an insertion sequence-dependent bias. Indeed, when we used the relative efficiencies observed in the “post-cloning pooling” experiment to correct the TAPE-1 unigram barcode frequencies measured in the “pre-cloning pooling” experiment, their correlation with the abundances of the corresponding pegRNAs in the plasmid pool improved (Pearson’s *r* = 0.87 → 0.94; **Supplementary Figure 1c**), and vice versa (Pearson’s *r* = 0.27 → 0.67; **Supplementary Figure 1e**).

## Screening additional monomers for their performance as DNA Ticker Tape

Our TAPE-1 construct exhibited sequential, directional editing, wherein the editing of any given site along the array was strongly dependent on all preceding sites having already been edited. This behavior is consistent with the DNA Ticker Tape design, as the key sequence must be inserted 5’ to any given monomer in order to complete the spacer that is recognized by any of the guide RNAs used. However, performance would presumably be corrupted by non-specific editing, *e.g.* if a guide were able to mediate edits to a non-write-head monomer despite several mismatches at the 5’ end of the spacer^19,20^.

Although TAPE-1 exhibited reasonable efficiency and specificity, we sought to explore whether this would be the case for other spacers. To this end, we designed and synthesized 48 TAPE constructs (TAPE-1 through TAPE-48), each derived from one of eight basal spacers that previously demonstrated reasonable efficiency in prime editing experiments^2,21,22^ and one of six design rules that vary monomer sequence, key sequence and key/monomer length (**Figure 2a,b**; **Supplementary Figure 2a**). In each of these 48 constructs, a 3xTAPE region was accompanied by a pegRNA-expressing cassette designed to target it with a 4-6 bp insertion (16 possible 2-bp barcodes followed by a 2-4 bp “key” sequence). We then transiently transfected HEK293T cells with PE2-encoding plasmid and a pool of 48 pegRNA-by-3xTAPE constructs and harvested them after four days. The 3xTAPE region was PCR-amplified from genomic DNA and sequenced.

**Figure 2.**
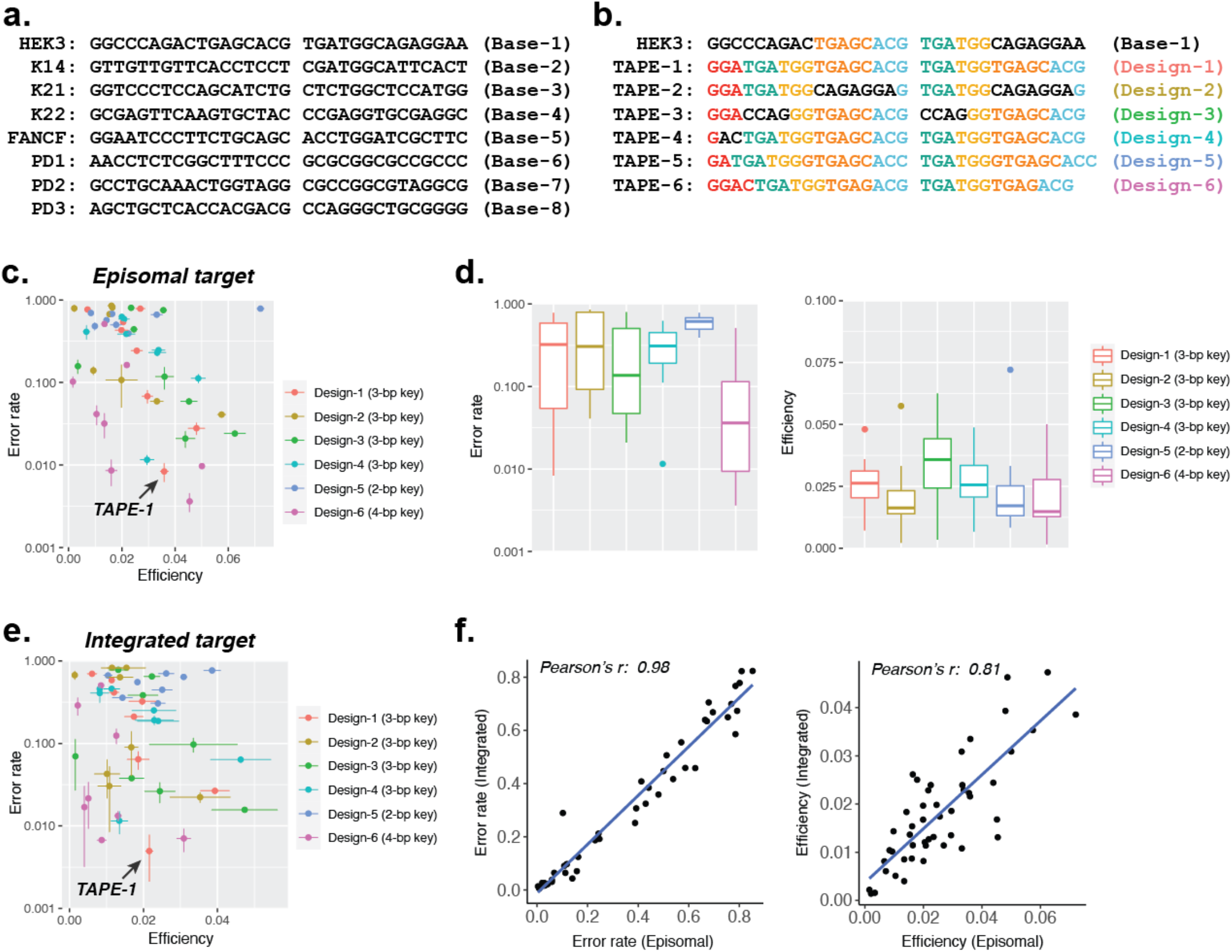
Characterizing diverse DNA Ticker Tape designs for efficiency and directional accuracy. **a.** Eight basal CRISPR spacers that previously demonstrated reasonable prime editing efficiencies^2,21,22^. **b.** Deriving TAPE designs (TAPE-1 to TAPE-6) from the basal HEK3 sequence via six distinct sequence shuffling procedures. The same procedures were also applied to the other basal targets to obtain the full set of 48 TAPE designs evaluated in this experiment (**Supplementary Figure 2**). **c.** Efficiency (fraction of edited reads out of all reads) vs. error rate (fraction of edited reads inconsistent with sequential, directional editing out of all edited reads) for 48 3xTAPE constructs on episomal DNA. Both horizontal and vertical error bars are standard deviations from 3 transfection replicates. **d.** Boxplots of the efficiencies and error rates of 3xTAPE constructs derived from 8 basal sequences for each of 6 design procedures. In general, a longer key sequence was associated with a lower error rate, while a longer insertion did not appreciably impact efficiency (*e.g.* NNGGAC with Design-6 vs. NNGA with Design-5). Boxplot elements represent: Thick horizontal lines, median; upper and lower box edges, first and third quartiles, respectively; whiskers, 1.5 times the interquartile range; circles, outliers. **e.** Same as panel **c**, but for constructs integrated via piggyBAC transposition. Both horizontal and vertical error bars are standard deviations from 3 transfection replicates. **f.** Correlation between the error rate (left) and editing efficiency (right) of each 3xTAPE construct either in the context of episomal DNA vs. integrated DNA.

We calculated two quantities for each 3xTAPE array: (1) efficiency, calculated by summing all edited reads and dividing by the total number of reads; and (2) error rate, calculated by summing all edited reads inconsistent with sequential, directional editing and dividing by the total number of edited reads (**Figure 2c**). Of note, our initial TAPE-1 construct had one of the lowest error rates among the 48 tested tapes. The only construct that had a lower error rate than TAPE-1 was TAPE-6, which was derived from the same basal spacer (HEK3) but had a 4-bp rather than 3-bp key sequence. Indeed, across the full experiment, a longer key sequence was associated with a lower error rate (**Figure 2d**). Performance differences between basal spacers were modest, with TAPEs based on the HEK3 and FANCF spacers exhibiting the best combination of efficiency and specificity (**Supplementary Figure 2b**). The efficiencies and error rates observed for individual monomers were highly consistent when we repeated the experiment with integration rather than transient transfection of the corresponding constructs (**Figure 2e,f**)

To understand how barcode length might affect tape efficiency and error rate, we focused on TAPE-1 and TAPE-27, which are respectively based on the HEK3 and FANCF spacers, both with 3-bp keys (**Supplementary Figure 3a**). We designed 5 pools of pegRNAs to insert 7-8 bp to TAPE-1 (4-5 bp barcode and 3-bp key), or 7-9 bp to TAPE-27 (4-6 bp barcode and 3-bp key). Each pegRNA pool was cloned and transfected separately, and analyses of efficiency and editing rate performed as previously. Editing efficiency ranged from 1.85% and 3.72%, with greater efficiency for longer TAPE-27 insertions (**Supplementary Figure 3b**). Error rates ranged from 0.91% to 1.95%, with lower error rates for longer insertions with both tapes (**Supplementary Figure 3c**). This result suggests that at least 6-bp barcodes can be used for DNA Ticker Tape, such that at least thousands (4^6 = 4096) of distinct signals can potentially be encoded and ordered.

## Recording and decoding of complex event histories

We next set out to ask whether we could apply DNA Ticker Tape to record, recover and decode complex event histories. We prepared a set of synthetic signals by individually cloning 16 individual pegRNA-expressing plasmids, each encoding a unique 2-bp barcode insertion to TAPE-1. We also prepared a polyclonal population of HEK293T cells with integrated 5xTAPE-1 to serve as the substrate for recording. Finally, we designed a set of five “transfection programs” -- complex event histories that we could attempt to record and then subsequently decode (**Figure 3a**).

**Figure 3.**
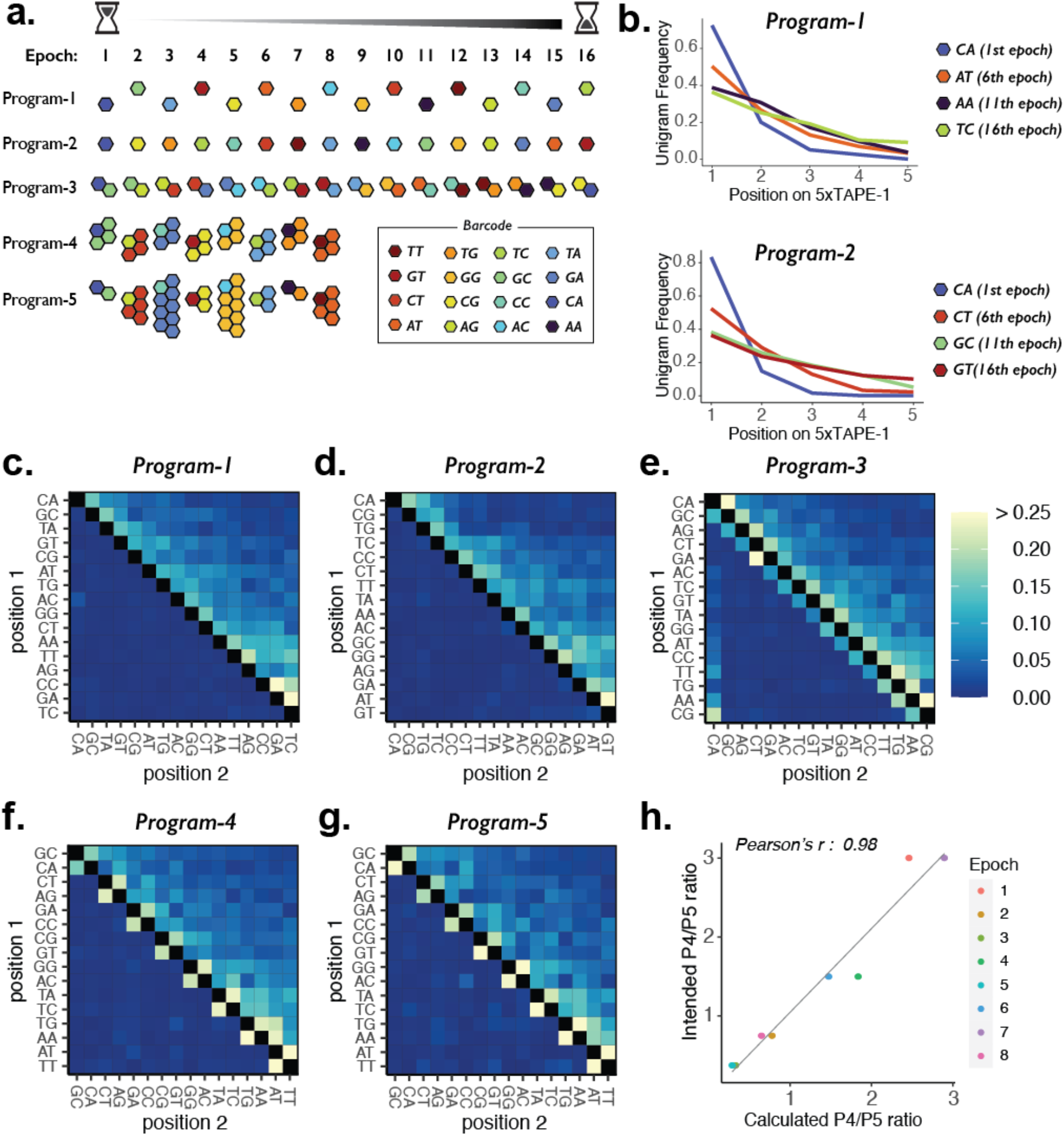
Transfection programs for 16 sequential signaling epochs. **a.** Schematic of five transfection programs over 8 or 16 epochs. For Programs 1 and 2, pegRNAs with single barcodes were introduced in each epoch for 16 epochs. The specific orders aimed to maximize (Program-1) or minimize (Program-2) the edit distances between temporally adjacent signals. For Program-3, pegRNAs with two different barcodes were introduced at a 1:1 ratio for 16 epochs, with one barcode always shared between adjacent epochs (and between epoch 1 and 16). For Programs 4 and 5, pegRNAs with two different barcodes were introduced either at constant ratio (1:3) or at varying ratios in each epoch (1:1, 1:2, 1:4, or 1:8) for 8 epochs, respectively. **b.** Barcode frequencies across 5 insertion sites in 5xTAPE-1 in Programs 1 and 2 following epoch 16. Barcodes introduced in early epochs are more frequently observed at the first site. **c-g.** Bigram transition matrix for Programs 1 (**c**), 2 (**d**), 3 (**e**), 4 (**f**), and 5 (**g**). Barcodes are ordered from early (left/top) to late (right/bottom). **h.** Calculated vs. intended relative frequencies between Programs 4 and 5.

At the beginning of each epoch of each transfection program, one or more synthetic signals was introduced to a population of HEK293T cells with integrated 5xTAPE-1 (5xTAPE-1(+) HEK293T) via transient transfection of plasmids expressing the corresponding pegRNA(s) and PE2. After each transfection, cells were passaged the next day into a new plate and excess cells were harvested for genomic DNA. 5xTAPE-1 from each epoch of each program was amplified and sequenced. Successive epochs occurred at 3-day intervals.

Programs 1 and 2 each consisted of a distinct, non-repeating sequence of the 16 synthetic signals, *i.e.* one per epoch. The specific orders aimed to maximize (Program-1) or minimize (Program-2) the edit distances between temporally adjacent signals. Based on sequencing of 5xTAPE-1 after Epoch-16, we observed that barcodes introduced in the early epochs were more frequent at the first target site (Site-1) than barcodes introduced at late epochs (**Figure 3b**). This is expected, as each editing round shifts more of the “write heads” to Site-2 (and subsequently to Site-3 to Site-5). A trivial decoding approach would be to simply arrange barcodes in the order of decreasing Site-1 unigram frequencies, but for both Programs 1 and 2, this results in an incorrect order (**Supplementary Table 2**).

However, the inference can be improved by leveraging the sequential aspect of DNA Ticker Tape, for instance by analyzing bigram frequencies. For example, if signal B preceded signal A, then we expect many more B-A bigrams than A-B bigrams at adjacent, edited sites within 5xTAPE-1. In **Figure 3c,d**, we show heatmaps of bigram frequencies, arranged by the true order in which the signals were introduced for Programs 1 and 2. Indeed, the bigram frequencies appear to capture event order information, evidenced by the gross excess of observations immediately above vs. immediately below the diagonal (*e.g.* in Program-1, CA-GC >> GC-CA). To leverage this information, we implemented the following algorithm: (1) initialize with the event order inferred from the Site-1 unigram frequencies; (2) iterate through adjacent epochs from beginning to end, and swap signals A and B if the bigram frequency of B-A is greater than A-B; (3) repeat step 2 until no additional swaps are necessary. For both Programs 1 and 2, this simple algorithm resulted in the correct ordering of the 16 signals, out of 16 factorial or 21 trillion possibilities (**Supplementary Table 2**).

The dearth of bigrams inconsistent with the true order, illustrated by the lack of signal below the diagonal in the Program-1 and Program-2 heatmaps (**Figure 3c,d**), indicates minimal interference between adjacent epochs, *i.e.* transfected pegRNAs from adjacent epochs did not overlap in their activities. To evaluate performance in the presence of such overlap, we designed Program-3, wherein two barcodes are introduced at each epoch, but adjacent epochs always share one barcode (**Figure 3a**). The concurrent transfection of two pegRNAs with distinct barcodes is evident in the resulting bigram frequency matrix, specifically by the signal both immediately above and below the diagonal (**Figure 3e**). Our aforedescribed decoding algorithm performs slightly worse on Program-3, with a single swap between epochs 4 and 5 required to revise the inferred order to the correct order (**Supplementary Table 2**).

Finally, we set out to ask whether the relative strength of signals could be inferred from DNA Ticker Tape. For this, we designed Programs 4 and 5, which share the same order of barcodes -- a pair in each epoch -- but with each pair at different ratios in the two programs. In Program-4, pegRNAs encoding each pair of barcodes were always mixed at a 1:3 ratio, whereas in Program-5, the same pairs for each epoch were mixed at a 1:1, 1:2, 1:4 or 1:8 ratio (**Figure 3a**). For both programs, the bigram frequency matrix is consistent with expectation, and the order of events was accurately inferred (**Figure 3f,g**; **Supplementary Table 3**). However, in addition, we were able to compare the relative ratios at which each pair of barcodes was introduced within each epoch between Programs 4 and 5, and found these to be well correlated with expectation (**Figure 3h**). Taken together, these results show that DNA Ticker Tape can record, recover and decode complex event histories including the order, overlap, and relative strength of signals.

## Recording and decoding short text messages

We next designed a strategy to record and decode short text messages to populations of cells with DNA Ticker Tape. In brief, we modified the Base64 binary-to-text encoding scheme by assigning each of the 64 possible 3-mers to 6-bit binaries. The Base64 scheme encodes uppercase and lowercase English characters, numbers from 0 to 9, and two symbols. In our TAPE64 scheme, we encoded uppercase English characters, four symbols and a whitespace, with two-fold or four-fold redundancy (**Figure 4a**).

**Figure 4.**
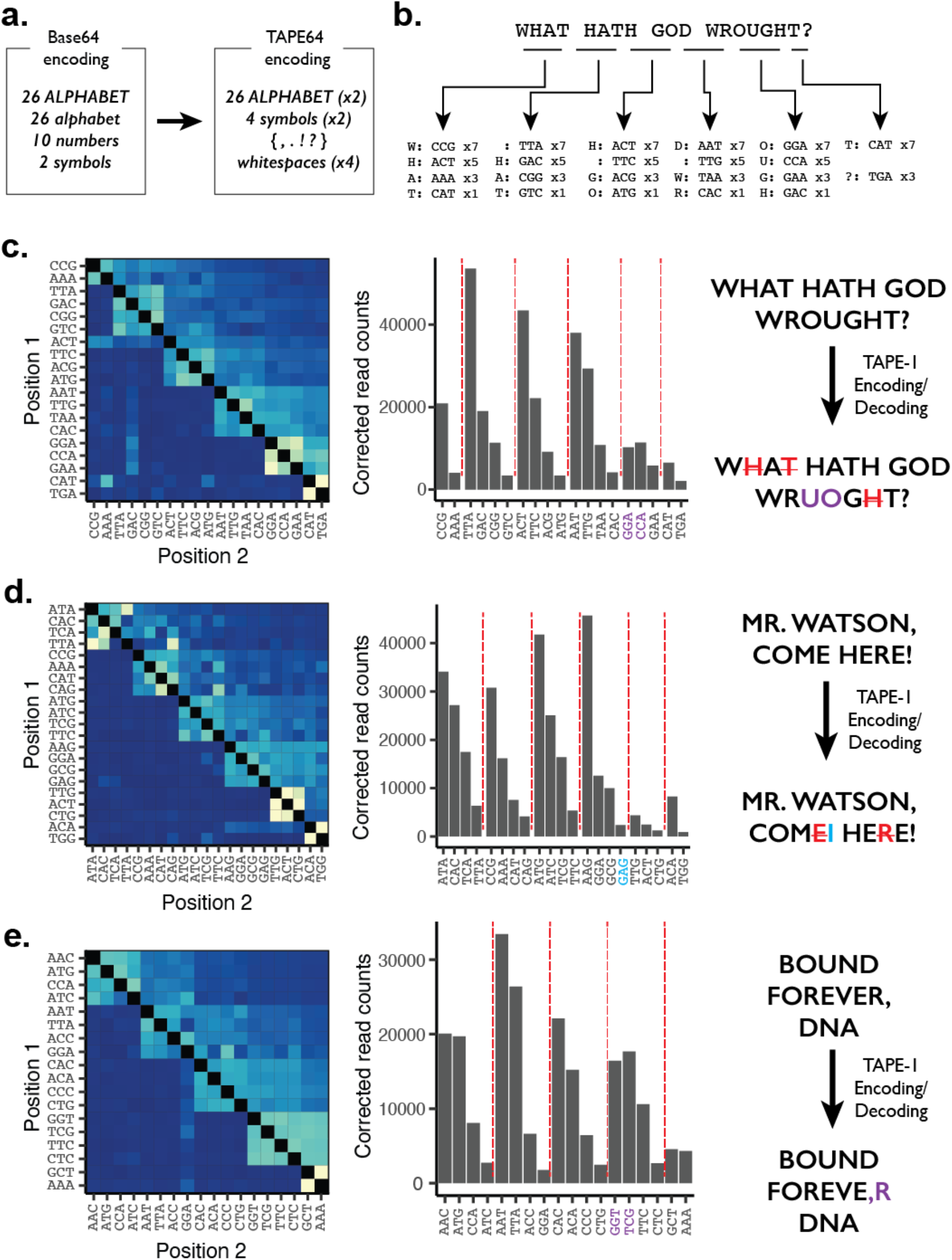
Recording and decoding short digital text messages with DNA Ticker Tape. **a.** Base64 binary-to-text was modified to assign 64 NNNGGA barcodes for TAPE-1 to 64 text characters. **b.** Illustration of the encoding strategy for “WHAT HATH GOD WROUGHT?”, which has 22 characters including whitespaces. The message is grouped into sets of 4 characters, converted to NNN barcodes according to the TAPE64 encoding table, and plasmids corresponding to each set mixed at a ratio of 7:5:3:1 for transfection. To encode 22 characters, we sequentially transfected 5 sets of 4 characters and 1 set of 2 characters 3 days apart to PE2(+) 5xTAPE-1(+) HEK293T cells. **c-e.** Decoding of 3 messages based on sequencing of 5xTAPE-1 arrays: (**c**) “WHAT HATH GOD WROUGHT?”, (**d**) “MR. WATSON, COME HERE!”, (**e**) “BOUND FOREVER, DNA”. For each message, the full set of NNNGGA insertions was first identified, and then co-transfected sets of characters identified from the bigram transition matrix (left). Within each set of characters inferred to have been co-transfected, ordering was based on corrected unigram counts (middle), resulting in the final decoded message (right). Misordered characters within each recovered message are colored purple, missing characters are colored red with strikethrough, and unintended characters are colored light blue.

We selected three messages to encode: (1) “WHAT HATH GOD WROUGHT?”, the first long-distance message transmitted by Morse code in 1844; (2) “MR. WATSON, COME HERE!”, the first message transmitted by telephone in 1876; and (3) “BOUND FOREVER, DNA”, a translation of a lyric from the 2017 song *DNA* by the K-pop music group *BTS*. Each message was split into sets of four characters. Plasmids encoding a given set of pegRNAs were concurrently transfected with a plasmid encoding PE2 to 5xTAPE-1(+) HEK293T cells at a ratio of 7:5:3:1, such that the ratio encoded the order of the four characters within each set (**Figure 4b**). As such, each full message could be recorded by five to six consecutive transfections spaced by three-day intervals.

To recover and decode the recorded messages, we harvested populations of cells corresponding to each message, and amplified and sequenced the tape region. From the resulting reads, we first identified all characters in the message by examining NNNGGA insertions at Site-1 of 5xTAPE-1. We then grouped these characters into sets by hierarchical clustering (**Supplementary Figure 4**), while also ordering these sets relative to one another, by applying the algorithm used for the previous experiment to the bigram transition matrix (**Figure 4c-e**). Finally, we arranged the four characters within each set by decreasing order of their editing score-corrected frequency, as within each set, earlier characters were encoded at a higher plasmid concentration.

For all three messages, our reconstructions of the original text were reasonable but imperfect. From the first message, 17/22 characters were correctly recovered and ordered, with three deletion errors and one swap between adjacent characters to yield “WA HATH GOD WRUOGT?” (**Figure 4c**). Of note, the deletion errors were due to repeated use of pegRNA barcodes ‘ACT’, ‘CAT’, and ‘GAC’ to encode multiple ‘H’ or ‘T’ characters, and as such were not expected to be recovered separately. From the second message, 20/22 characters were correctly recovered and ordered, with two deletions and one insertion to yield “MR. WATSON, COMI HEE!” (**Figure 4d**). From the third message, 16/18 characters were correctly recovered and ordered, with a single swap between adjacent characters to yield “BOUND FOREVE,R DNA” (**Figure 4e**). Despite these errors, our experiment demonstrates the potential of DNA Ticker Tape to digitally record the content and order of information to the genomes of populations of mammalian cells.

## Ordered recording and decoding of cell lineage histories

Beginning with <u>G</u>enome <u>E</u>diting of <u>S</u>ynthetic <u>T</u>arget <u>A</u>rrays for <u>L</u>ineage <u>T</u>racing (GESTALT), a number of approaches have been developed that leverage stochastic *in vivo* genome editing to generate a combinatorial diversity of mutations that irreversibly accumulate during development to a compact DNA barcode^6,23^. Such stochastically evolving barcodes mark cells and enable inference of their developmental lineage relationships based on patterns of shared mutations. However, despite their promise, unordered lineage recorders remain sharply limited by several technical challenges, including: (1) a failure to explicitly record the order of editing events, which renders phylogenetic reconstruction of cell lineage highly challenging^24,25^; (2) a reliance on double-stranded breaks (DSBs) and nonhomologous end-joining (NHEJ) to introduce edits; DSBs are cytotoxic and frequently delete consecutively located targets within a barcode; and (3) the number of target sites available to CRISPR-Cas9 decreases as sites are irreversibly edited, which makes it challenging to sustain a continuous rate of lineage recording throughout development.

The ordered nature of DNA Ticker Tape, its reliance on prime editing, and the fact that only the “write head” monomer is active at any given moment, potentially address all of these limitations at once. To explore the utility of DNA Ticker Tape to simplify genome editing-based cell lineage tracing, we performed an experiment to record early cell divisions of a monoclonal cell line. For this, we constructed a HEK293T cell line that expresses doxycycline (Dox)-inducible PE2 (iPE2(+) HEK293T). We also designed and cloned a lentiviral construct that includes: (1) 5xTAPE-1 sequence, associated with a random 8-bp barcode region (TargetBC) at its 5’-end; and (2) a constitutive pegRNA expression cassette that targets TAPE-1 for a 6-bp insertion (<u>NNN</u>GGA; <u>NNN</u> is subsequently referred to as InsertBC; GGA is the “key” sequence for TAPE-1) (**Figure 5a**). Lentiviral transduction at a high multiplicity of infection (MOI) was followed by serial dilution to isolate a monoclonal cell line that grew from a single cell to ~200,000 cells via ~18 doublings over 32 days in the presence of Dox (**Figure 5b**).

**Figure 5.**
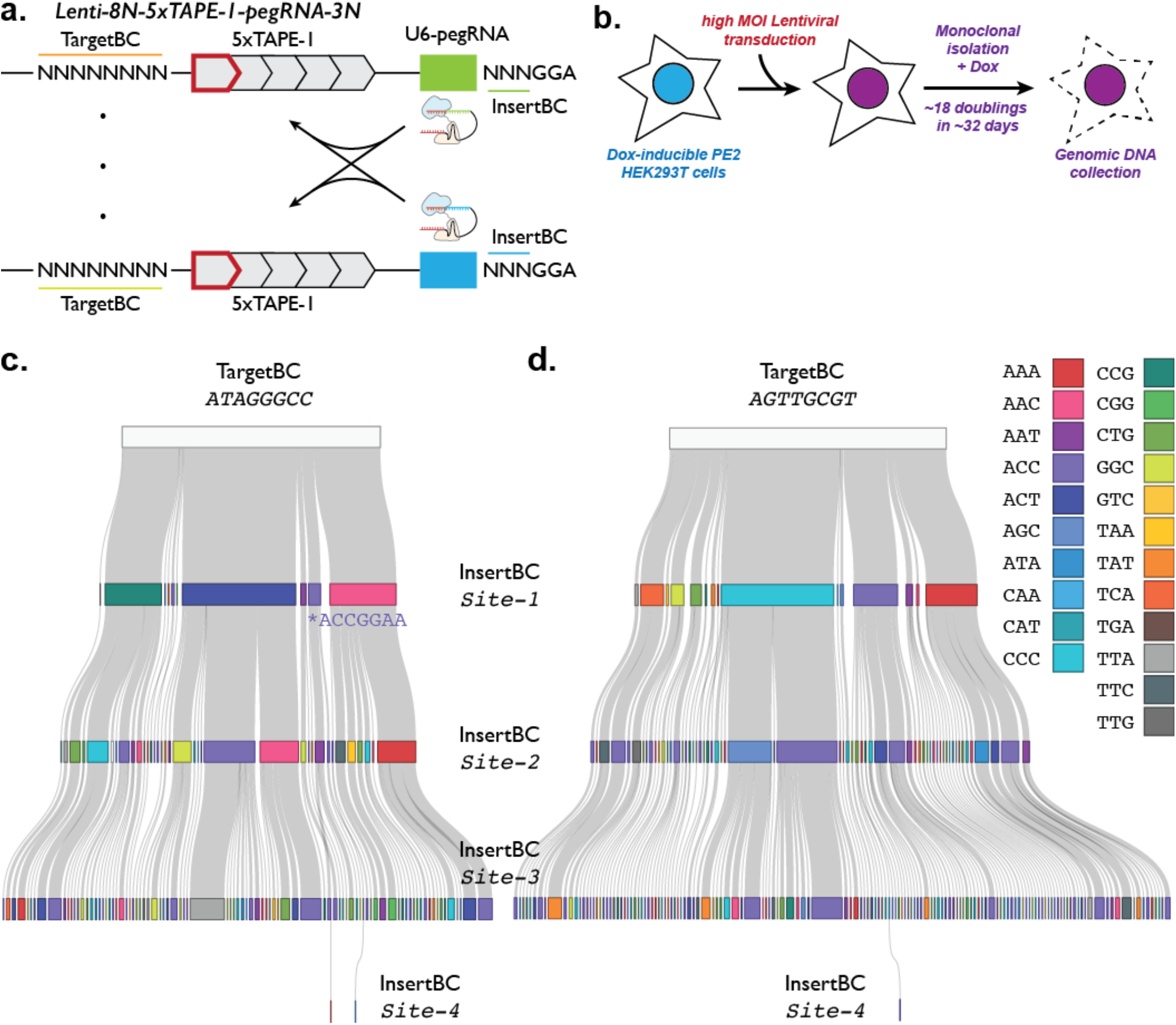
Reconstruction of early cell division events using 5xTAPE-1 editing patterns. **a.** Description of lentiviral vector used in the DNA Ticker Tape-based lineage tracing experiment. The lentiviral integration region includes a 5xTAPE-1 sequence associated with an 8-bp random barcode (TargetBC), and a pegRNA expression cassette. pegRNA targets TAPE-1 and inserts 6-bp, where the first 3-bp is the random barcode (InsertBC) and the last 3-bp is the “key” sequence of GGA for TAPE-1. **b.** Schematic of lineage tracing experiment in a monoclonal HEK293T cell line. During its expansion, 23 pegRNAs expressed by 23 TargetBC-defined integrants, collectively encoding 22 unique InsertBCs (one recurrent by chance), compete to mediate insertions at the “write heads” of 23 TAPE arrays. **c-d.** Reconstruction of editing events at TargetBC (ATAGGGCC) (**c**) and TargetBC (AGTTGCGT) (**d**). A prime editing error resulting in a 7-bp insertion (<u>ACC</u>GGA<u>A</u>) at Site-1 of the TargetBC (ATAGGGCC) TAPE array is noted with an asterisk. Each insertion is colored by the 3-bp identity following the key on the right.

The region containing the TargetBC and 5xTAPE-1 was then amplified from genomic DNA and sequenced. To enhance data quality, we used unique molecular identifiers (UMIs) appended by a linear amplification phase prior to PCR, as well as paired-end sequencing to bidirectionally cover the amplicon. After UMI deduplication, we combined reads derived from 16 PCR replicates and, based on read count frequencies, identified 23 TargetBC barcodes, each marking a unique insertion of the lentiviral construct into the genome of the monoclonal cell line (**Supplementary Figure 5a**). Correspondingly, although there are 4^3^ or 64 potential InsertBC barcodes, only 22 were observed at appreciable frequencies, suggesting that one recurred by chance, possibly <u>ACC</u> based on its considerably greater usage (**Supplementary Figure 5b**). For other InsertBC barcodes, the observed variation in usage is probably explained by some combination of: (1) variation in pegRNA expression levels due to site-of-integration effects; and (2) differences in insertional efficiency. Based on the frequencies of usage of InsertBC barcodes, we calculated their Shannon entropy to be 3.64 and 3.70 bits for Site-1 and Site-2, respectively, out of a theoretical maximum of 4.46 bits if the probabilities of the 22 InsertBC barcodes were equal. The Kullback-Leibler divergence of InsertBC frequencies of Site-2 from Site-1 (*D*_*KL*_*(Site-2||Site-1)*) was 0.13 bits, suggesting InsertBC barcode usage is similar between TAPE-1 sites.

Across all 23 TargetBC-defined tape integrations, we identified a total of 6644 unique “alleles”, each present at a frequency >1% of that of the most common allele at the same integration site (**Supplementary Table 4**). In contrast with the edits obtained with unordered lineage recorders, the ordered insertional barcodes of DNA Ticker Tape are straightforward to organize into cell lineage trees (**Figure 5c-d**; **Supplementary Figure 6**). For example, TargetBC (ATAGGGCC) was highly edited at its first three sites (99.9% edited at Site-1; 97.7% edited at Sites 1-3). A total of 90 alleles can be organized into 10 groups based on their Site-1 insertions, which further branch to 36 groups based on their Site-2 insertions, and finally into the 90 groups based on their Site-3 insertions (**Figure 5c**). As far as why editing did not extend beyond Site-3, we observed a 1-bp deletion near the PAM sequence of Site-4 of nearly all TargetBC (ATAGGGCC) reads, presumably inactivating it altogether for this particular integrant. Interestingly, we rarely observe the editing of the fourth TAPE-1 site along with a deletion of the fifth TAPE-1 monomer (2 alleles; 0.3% of all TargetBC (ATAGGGCC) reads), suggesting that occasional contraction of the monomer array by one unit resulted in a corrected and editable Site-4 sequence.

Prime editing errors are rare, but do occur. For example, among the insertions at Site-1 of TargetBC (ATAGGGCC), we observe <u>ACC</u>GGA (4 alleles; 1.0% of reads) but also <u>ACC</u>GGA<u>A</u> (8 alleles; 4.7% of reads) (**Figure 5c**, **Supplementary Table 4**). The latter, unexpected 7-bp insertion only occurs at Site-1 of TargetBC (ATAGGGCC), and not downstream sites of this tape integrant nor in any site of other tape integrant alleles. This pattern supports the shared ancestry of all alleles bearing this insertion at Site-1, presumably a stochastic prime editing error that occurred once during the early proliferation of this monoclonal line.

For the programmed insertions (*i.e.* the 22 x 3-bp barcodes), how confident can we be that a given insertion at a given site, shared across alleles, is due to a single event in a common ancestor, rather than recurrence? Focusing on TargetBC (ATAGGGCC), we note that the distribution of InsertBCs at Site-1 of TargetBC (ATAGGGCC) is substantially different from the overall InsertBC distribution, presumably reflecting early, stochastic events (**Figure 5c**; **Supplementary Figure 5d**). To quantify this, while the Shannon entropy of insertion events at Site-1 of TargetBC (ATAGGGCC) is 1.93 bits, the Kullback-Leibler divergence of these insertions vs. Site-1 insertions across all integrants (*D*_*KL*_*(Site-1*_*ATAGGGCC*_*||Site-1*_*All*_)) is 1.70 bits. This is consistent with alleles sharing the same InsertBC at Site-1 generally sharing a common ancestor. Although the rate of editing is somewhat too slow to construct an unequivocal, binary tree, we can infer that the most frequent insertion event at Site-1 (ACTGGA or 1.ACT; 44% of reads) probably occurred after the first or second cell division. Most 1.ACT alleles further branch to 1.ACT-2.ACC and 1.ACT-2.AAC at Site-2 (respectively 45% and 34% of 1.ACT reads), insertions which probably occurred between the second and fourth cell divisions.

Looking further down this tree, if we consider <u>all</u> insertions at Site-2 or Site-3, irrespective of their Site-1 state, the Kullback-Leibler divergence of InsertBCs vs. the background distribution is substantially lower than for Site-1 (*D*_*KL*_*(Site-2*_*ATAGGGCC*_*||Site-1*_*All*_) = 0.67 bits; *D*_*KL*_*(Site-3*_*ATAGGGCC*_*||Site-1*_*All*_) = 0.55 bits). This is expected, as the number of cells is presumably much larger as these sites are being edited, and in this calculation we are considering all barcodes including instances where a given insertion may have occurred recurrently (*i.e.* “identity by state”). However, within any given subtree, as defined by insertions at earlier site(s) along the tape, the Kullback-Leibler divergence to the background distribution remains high (mean *D*_*KL*_(*Site-2*_*ATAGGGCC*_ subtree|*|Site-1*_*All*_) = 2.72 bits; mean *D*_*KL*_*(Site-3*_*ATAGGGCC*_subtree|*|Site-1*_*All*_) = 2.77 bits). As such, conditional upon earlier edits along the tape, recurrent observations of a given insertion within a subtree likely correspond to single events (*i.e.* “identity by descent”).

Lineage trees for other TargetBC-defined tape integrants are similarly straightforward to construct. For example, TargetBC (AGTTGCGT) has 162 alleles (**Figure 5d**). Nearly all of these alleles are collapsed to 3 monomers and contain an error at the initiating key (GGA → GGAA), alterations inferred to have been present at this integrant in the founding monoclonal cell, potentially introduced during oligo synthesis, cloning or lentiviral production. Surprisingly, despite the single-base insertion at the key sequence, this tape was still functional, with over 99% of Site-1 reads edited. The Shannon entropy of Site-1 of this tape is 2.62 bits, which diverges 1.40 bits from Site-1 insertions across all integrants (*D*_*KL*_*(Site-1*_*AGTTGCGT*_*||Site-1*_*All*_)) (**Figure 5d**).

Additional examples are shown in **Supplementary Figure 6**. Consistent with our first DNA Ticker Tape experiment (see earlier discussion of “pseudo-processivity”), we see evidence for site-of-integration effects on the rate of tape editing. For example, the tree shown in **Supplementary Figure 6a** illustrates a TargetBC with relatively fast editing, while the tree shown in **Supplementary Figure 6b** illustrates a TargetBC with relatively slow editing. Whereas “fast tape” brings us closer to an unequivocal, binary tree, the validity of trees obtained with “slow tape” may be compromised by recurrences of the same edit within subtrees (**Supplementary Table 5**). Overall, these results demonstrate how DNA Ticker Tape can be used to reconstruct cell lineage histories without a need for specialized computational methods to infer the order of editing events.

## Editing and recovering longer TAPE arrays

The information density and maximum potential recording duration of DNA Ticker Tape are directly proportional to the number of consecutive TAPE monomers. However, long tandem arrays can be challenging to work with from a technical perspective, *e.g.* difficult to synthesize, clone and maintain; prone to instability during *in vivo* DNA replication or repair as well as during *in vitro* PCR; difficult to accurately and fully sequence.

To evaluate the extent to which such issues might be limiting in practice, we attempted to create a synthetic minisatellite in the form of 12 or 20 repeats of the 14-bp TAPE-1 monomer (12xTAPE-1 and 20xTAPE-1). 12xTAPE-1 was synthesized as single-stranded DNA (IDT) and 20xTAPE-1 as a plasmid (GenScript). PCR amplicons of each array were cloned into the piggyBAC vector via Gibson assembly. Of note, cloned constructs were used “as-is”, even though it is possible that some degree of variation in repeat number was already present (**Supplementary Figure 7**). We then integrated piggyBAC vectors bearing ~12xTAPE-1 or ~20xTAPE-1 into HEK293T cells expressing both PE2 and pegRNAs targeting TAPE-1 for NNNGGA insertions (PE2(+) 3N-TAPE-1-pegRNA(+) HEK293T) in triplicate. We cultured these cell lines for 40 days before collecting genomic DNA. PCR amplification of TAPE-1 was followed by standard library construction and sequencing on the Pacific Bioscience Sequel platform to obtain circular consensus sequencing (CCS) reads.

We grouped CCS reads within each replicate based on a degenerate 8-bp barcode (TargetBC), as these presumably derived from the same integration. On average, each TargetBC group had 3.1 ± 3.4 and 3.8 ± 5.7 reads for ~12xTAPE-1 and ~20xTAPE-1, respectively. Although some variability in repeat length was observed within many TargetBC-defined groups, arrays longer than the intended 12x or 20x lengths were rare. As this suggests contraction from the original array length to be more likely than expansion, we selected the largest array within each TargetBC group as representative. Of representative CCS reads for 4784 and 6254 integrated arrays for 12xTAPE-1 and 20xTAPE-1, respectively, the overwhelming majority (>99.5%) exhibited clear patterns of sequential, directed editing (**Supplementary Figure 8**). Although ~20xTAPE-1 arrays tended to be longer, the extent of sequential editing between the two conditions was very similar (mean of 3.9 ± 1.9 events and 4.1 ± 2.1 events for ~12xTAPE-1 and ~20xTAPE-1, respectively). This is consistent with the supposition that because DNA Ticker Tapes have only one active “write head”, the rates at which they are written to should be independent of their length. As with previous experiments, we observe extensive variation in editing rate across TargetBCs, which we again suspect is due to site-of-integration effects. In terms of the maximum extent to which any given tape was edited, we observed one TargetBC for which 14 distinct 3-bp insertion events were recorded along on a 14xTAPE-1.

This experiment illustrates that it is possible to construct and use synthetic minisatellites corresponding to at least 20 monomers as DNA Ticker Tape, and that sequential recording of at least 14 consecutive events is possible. Nonetheless, further experiments are required to quantify the extent to which variation in synthetic minisatellite length is due to: (1) piggyBAC vector heterogeneity, *i.e.* variation that existed prior to integration; (2) DNA replication and microsatellite instability in HEK293T cells; (3) DNA repair subsequent to prime editing-induced nicks; and/or (4) PCR amplification artifacts. Of note, the observed variation in array length tends to occur within the unedited portion of the tape (**Supplementary Figure 8**). We have yet to observe any clear examples of “information erasure”, possibly because the edits themselves disrupt the tandem repeats, inhibiting processes that might otherwise lead to erasure from spreading proximal to the “write head”.

## Discussion

Digital systems represent information through both the content and order of discrete symbols, with each symbol drawn from a finite set. Digital systems are ancient, and include written text, morse code, binary data, and, of course, DNA itself. In this proof-of-concept of DNA Ticker Tape, we demonstrate how sequential genome editing of a monomeric array constitutes an artificial digital system that is operational within living eukaryotic cells, capable of “writing” thousands of discrete symbols to DNA in an ordered fashion.

How might DNA Ticker Tape be used? Specific symbols, *i.e.* insertional barcodes, are mediated by specific pegRNAs. Although we have not focused on it here, we and other groups have developed various approaches to couple biological signals to genome editing. For example, several groups have engineered guide RNAs whose activity is dependent on the binding of specific small molecules or ligands^26–28^. In parallel to this work, we developed a prime editing-based system for signal-specific recording called ENGRAM, wherein biological signals of interest are coupled to the Pol-2-mediated transcription of specific guide RNAs, whose expression then programs the insertion of signal-specific barcodes to a DNA Tape. At least in principle, ENGRAM is compatible with DNA Ticker Tape, potentially enabling the temporal dynamics of multiple biological signals or other cellular events to be recorded and resolved.

Indeed, in principle, the identity and order of thousands of distinct biological signals can be recorded simultaneously with DNA Ticker Tape by coupling each signal to the expression or activity of a pegRNA that mediates a specific insertional barcode. Recording channels could be further partitioned via multiple TAPE arrays composed of different monomers, each with its own speed and error rate characteristics, with minimal interference between them. The information density of DNA Ticker Tape is predicted to scale exponentially with any improvements in prime editing’s ability to make longer insertions efficiently and precisely.

However, even with the stochastic recording (*i.e.* editing that is not coupled to any specific biological signal), the ability to temporally order events is immediately useful for developmental lineage tracing. In a proof-of-concept experiment, we show how DNA Ticker Tape overcomes the major limitations of earlier editing-based lineage recorders like GESTALT, most critically by eliminating ambiguity about the order in which editing events occurred, but also by minimizing the risk of inter-target deletion and ensuring a constant number of active targets over time.

What are the limits of this approach? It is plausible to imagine that with an array of 100 TAPE monomers and an editing rate of ~2 insertions per cell division, one could reconstruct a largely unequivocal binary tree over 50 generations, more than sufficient for Sulston-like reconstructions^29^ of mouse or zebrafish development. We further envision that a single, synthetic DNA construct that encoded a prime editing enzyme, a large monomeric recording array, and a combination of stochastic and signal-specific pegRNAs, could be used to simultaneously record both lineage and biological signals during the development of a multicellular organism. Because it leverages prime editing, such recording would be expected to continue to take place in non-mitotic cells such as neurons^2^, capturing signals longitudinally. Although it may be challenging to engineer, such an artificial recorder device would allow us to take full advantage of DNA as an *in vivo* digital recording medium, richly capturing the breadth and order of signals and events throughout metazoan development, as well as in other biological systems of interest.

## Acknowledgements

We thank the members of the Shendure Lab, as well as members of the Allen Discovery Center for Cell Lineage Tracing, for helpful discussions. We thank David Liu’s lab for sharing the prime editing plasmids. This work was supported by a grant from the Paul G. Allen Frontiers Group (Allen Discovery Center for Cell Lineage Tracing to J.S.) and the National Human Genome Research Institute (UM1HG011586 to J.S.). J.C. is a Howard Hughes Medical Institute Fellow of the Damon Runyon Cancer Research Foundation (DRG-2403-20). J.S. is an Investigator of the Howard Hughes Medical Institute.

## Data availability statement

Raw sequencing data have been uploaded on Sequencing Read Archive (SRA) with associated BioProject ID PRJNA757179. Plasmids encoding DNA Ticker Tape construct (piggyBAC-5xTAPE-1-BlastR) and pegRNA (pU6-CApegTAPE1) have been deposited to Addgene (ID 175808 and 175809).

## Code availability statement

Custom analysis codes for this project are available at Github: https://github.com/shendurelab/DNATickerTape

## Materials and Methods

### Plasmid cloning

Both pegRNA and TAPE constructs were cloned either using Gibson assembly (Gibson Assembly Master Mix, New England Biolab) or ligation after restriction (T4 DNA Ligase, New England Biolabs). For the Gibson assembly protocol, inserts of interest, usually ordered in the form of single-stranded DNA (IDT; Ultramer, up to 200-bp, or IDT oPool, up to 350-bp), were amplified using polymerase chain reaction (PCR; KAPA HiFi polymerase) and converted into double-stranded DNA molecules. For ligation, single-stranded DNAs (IDT) were annealed to have 4 bp overhangs in both ends of double-stranded DNAs, which is a substrate for T4 DNA ligase. Cloning backbones were digested either with BsaI-HFv2 or BsmBI-v2 (NEB), gel-purified, and mixed with inserts in the Gibson Assembly reaction. A small amount (1-2 uL) of Gibson Assembly reaction mix or T4 ligation mix was added to NEB Stbl cell (C3040) for transformation, and grown at 30°C or 37°C for the plasmid DNA preparation (Qiagen miniprep). The resulting plasmids were sequence-verified using Sanger sequencing (Genewiz).

### Tissue culture, transfection, lentiviral transduction, and transgene integration

The HEK293T cell line was purchased from ATCC, and maintained by following the recommended protocol from the vendor. HEK293T cells were cultured in Dulbecco’s modified Eagle’s medium with high glucose (GIBCO), supplemented with 10% fetal bovine serum (Rocky Mountain Biologicals) and 1% penicillin-streptomycin (GIBCO). Cells were grown with 5% CO2 at 37°C.

For transient transfection, HEK293T cells were cultured to 70-90% confluency in a 24-well plate. For prime editing, 375 ng of Prime Editor-2 enzyme plasmid (Addgene #132776) and 125 ng of pegRNA plasmid were mixed and prepared with a transfection reagent (Lipofectamine 3000) following the recommended protocol from the vendor. Cells were cultured for four to five days after the initial transfection unless noted otherwise, and its genomic DNA was harvested following cell lysis and protease protocol from Anzalone et al.^2^.

For lentivirus generation, about 300,000 HEK293T cells were seeded to each well in a 6-well plate and cultured to 70-90% confluency. The lentiviral plasmid was transfected along with the ViraPower lentiviral expression system (ThermoFisher), following the recommended protocol from the vendor. Lentivirus was harvested following the same protocol, concentrated overnight using Peg-it Virus Precipitation Solution (SBI), and used within 1-2 days to transduce either K562 or HEK293T cells without a freeze-thaw cycle. To achieve high multiplicity of infection, we used MagnetoFection protocol (OZ Bioscience). For the lineage-tracing experiments, transduced cells were serially diluted and seeded to 96-well plates to identify monoclonal lines. Dox concentrations were maintained by having 10 mg/L in the initial culture and replenished every five days considering 24 to 48 half-life of Dox in culturing media.

For transposase integration, 500 ng of cargo plasmid and 100 ng of Super piggyBAC transposase expression vector (SBI) were mixed and prepared with a transfection reagent (Lipofectamine 3000) following the recommended protocol from the vendor and transfected to a confluent 24-wells. The monoclonal Dox-inducible PE2 cell line was generated by integrating PE2 using the piggyBAC transposase system and selecting clones by prime-editing activity, as previously described^22^.

### Genomic DNA collection and sequencing library preparation

The targeted region from collected genomic DNA was amplified using two-step PCR and sequenced using Illumina sequencing platform (NextSeq or MiSeq). The first PCR reaction (KAPA Robust polymerase) included 1.5 uL of cell lysate, 0.04 to 0.4 uM of forward and reverse primers in a final reaction volume of 25 uL. We programmed the first PCR reaction to be: (1) 3 minutes at 95°C, (2) 15 seconds at 95°C, (3) 10 seconds at 65°C, (4) 90 seconds at 72°C, (5) 25-28 cycles of repeating step 2 through 4, and (6) 1 minute at 72°C. Primers included sequencing adapters to their 3′-ends, appending them to both termini of PCR products that amplified genomic DNA. After the first PCR step, products were assessed on 6% TBE-gel and purified using 1.0X AMPure (Beckman Coulter) and added to the second PCR reaction that appended dual sample indexes and flow cell adapters. The second PCR reaction program was identical to the first PCR program except we ran it for only 5-10 cycles. Products were again purified using AMPure and assessed on the TapeStation (Agilent) before being denatured for the sequencing run.

For appending 10-bp unique molecular identifiers (UMI), we performed the PCR reaction in three steps: First, genomic DNA was linearly amplified in the presence of 0.04 to 0.4 uM of single forward primer in two PCR cycles using KAPA Robust polymerase. Specifically, we programmed the UMI-appending linear PCR reaction to be: (1) 3 minutes and 15 seconds at 95°C, (2) 1 minute at 65°C, (3) 2 minutes at 72°C, (4) 5 cycles of repeating step 2 and 3, (5) 15 seconds at 95°C, (6) 1 minute at 65°C, (7) 2 minutes at 72°C, and (8) another 5 cycles of repeating step 6 and 7. Second, this reaction was cleaned up using 1.5X AMPure, and then to a second PCR with forward and reverse primers: (1) 3 minutes at 95°C, (2) 15 seconds at 95°C, (3) 10 seconds at 65°C, (4) 90 seconds at 72°C, (5) 25-28 cycles of repeating step 2 through 4, and (6) 1 minute at 72°C. In this case, the forward primer binds upstream of the UMI sequence and is not specific to the genomic locus. Finally, after PCR amplification, products were cleaned up using AMPure magnetic beads (1.0X, following the protocol from Beckman Coulter) and added to the third and last PCR reaction that appended dual sample indexes and flow cell adapters. The run parameters for the third PCR reaction was the same as the second PCR reaction, except only 5-10 cycles of repeating step 2 through 4 were used. TAPE construct sequences and PCR primer sequences are listed in **Supplementary Table 6** and **Supplementary Table 7**, respectively.

For long-read amplicon sequencing library preparation, we used a two-step PCR protocol: the first PCR reaction (KAPA Robust polymerase) included 1.5 uL of cell lysate, 0.04 to 0.4 uM of forward and reverse primers in a final reaction volume of 25 uL. We programmed the first PCR reaction to be: (1) 3 minutes at 95°C, (2) 15 seconds at 95°C, (3) 10 seconds at 65°C, (4) 3 minutes at 72°C, (5) 25-28 cycles of repeating step 2 through 4, and (6) 1 minute at 72°C. After the first PCR step, products were assessed on 6% TBE-gel and purified using 0.6X AMPure (Beckman Coulter) and added to the second PCR reaction that appended PacBio sample indexes. The second PCR reaction program was identical to the first PCR program except we ran it for only 5-10 cycles. Products were again purified using AMPure and assessed on the TapeStation (Agilent) and sequenced on Sequel (Pacific Biosciences; Laboratory of Biotechnology and Bioanalysis, Washington State University).

### Sequencing data processing and analysis

For sequencing experiments shown in **Figures 1** and **Figure 2** (and also shown in **Supplementary Figures 1-3**), sequencing libraries were single-end sequenced to cover the TAPE arrays from one direction. For recording experiments shown in **Figure 3** and **Figure 5** (and also shown in **Supplementary Figures 4 to 6**), sequencing libraries were paired-end sequenced to cover the entire array from both directions. Paired reads were then merged using PEAR^30^ with default parameters to reduce sequencing errors. Insertion sequences, either in the form of NNGGA 5-mer to NNNGGA 6-mer were extracted from sequencing reads of the TAPE arrays, including 2xTAPE-1, 3xTAPE-1, and 5xTAPE-1, using pattern-matching software such as Regular Expression (package *REGEX*) in Python. Insertions (4 to 6 bp) on 3xTAPE-1 to 3xTAPE-48 were also extracted using *REGEX* pattern-matching software.

In the sequential signaling epochs experiment shown in **Figure 3**, we first extracted 5-mer insertions from the 5xTAPE-1 sequencing reads, and used a k-means clustering algorithm to filter out possible PCR/sequencing errors with low read counts. Such filtering removed all reads that have the wrong Key sequence (GGA in the case of TAPE-1), leaving a set of 16 possible 5-mer sequences in the form of NNGGA. Across 5 repeats of insertion sites on 5xTAPE-1, we calculated the separate unigram frequencies in each site, which was used to build the Unigram order as shown in **Supplementary Tables 2-3**. Bigram frequencies between the adjacent insertion sites (Site-1 and Site-2, Site-2 and Site-3, Site-3 and Site-4, and Site-4 and Site-5 pairs) were combined, normalized across row and column, and used to build the bigram transition matrices as shown in **Figure 3c-g**. For ordering the barcodes according to their transfection history, we first generated a Unigram order by sorting its relative frequency on Site-1, where barcodes were assumed to have transfected earlier if they appeared more frequently in Site-1 than other sites. Using the resulting Unigram order as the initial order, we implemented an iterative algorithm where we pass through the order, from early to late, swap the order if their bigram frequency is inconsistent with the order, and restart the pass unless there have been no swaps in a single pass.

In the short digital text encoding experiment shown in **Figure 4**, we extracted 6-mer insertions, corrected the read-counts of each 6-mers by their editing efficiencies (using separately measured insertion frequency and respective plasmid abundance, similar to described in **Supplementary Figure 1c,e**), used a k-means clustering algorithm to identify NNNGGA barcodes, and built the bigram transition matrix as described in the paragraph above. We first analyzed the bigram transition matrices using a hierarchical clustering algorithm, with default parameters given in the *R* software (using Euclidean distance measure and complete-linkage clustering method, as described in **Supplementary Figure 4**). Putative sets of barcodes (co-transfection sets with generally 2-4 barcodes) were visually identified based on the dendrogram, and used to group barcodes in the output bigram order of the algorithm used above. The order within the co-transfection sets was determined using the corrected unigram counts combined across all five sites, where more abundant barcodes were assigned to be earlier within the set. Barcodes were mapped back to the text following the encoding table (**Supplementary Table 8**)

In the lineage-tracing experiments, an extra linear PCR step was added to append a 10-bp unique molecular identifier (amplicon-UMI) to each amplicon (TargetBC-5xTAPE-1, where an 8-bp TargetBC sequence identifies the lentiviral integrant), before the exponential PCR amplification in 16 PCR-replicates. To minimize the sequencing errors, we used paired-end sequencing to bidirectionally cover the TargetBC-5xTAPE-1 amplicon, then merged using *PEAR*, as described above. To further minimize PCR errors, sequencing reads that share the same amplicon-UMI were collapsed into a single most frequent read, and any amplicon-UMI sequences that appeared less than 3 times within the PCR replicate were excluded. Combining all 16 PCR-replicates after amplicon-UMI collapsing and filtering resulted in ~370k total reads. These reads were further collapsed based on the shared TargetBC-5xTAPE-1 sequence, and used to identify the set of TargetBCs and InsertBCs as shown in **Supplementary Figure 5**. Assuming that the observation probability of erroneous reads (with PCR and/or sequencing errors) correlates with the frequency of correct, basal reads, we set the threshold of 1% of the largest read counts within the set of TargetBC-5xTAPE-1 reads sharing the same TargetBC to exclude reads with PCR and/or sequencing errors, resulting in 6644 alleles and counts for each allele (**Supplementary Table 4**). Insertion sequences or InsertBCs were identified using text pattern matching. Possible mutations on the TAPE-1 sequence were visually inspected to avoid incorrect assignment of InsertBCs in one of the possible sites within the 5xTAPE-1 array (with or without possible contractions or expansions).

In the long-read sequencing experiment, 12xTAPE-1 and 20xTAPE-1 sequences were isolated from Pacific Biosciences circular consensus (CCS) reads. The number of TAPE-monomers and insertions were calculated using sequential text-matching around insertions and the expected length of the array based on insertion counts. Reads without a match between expected length and observed length were filtered out. Each 12xTAPE-1 and 20xTAPE-1 construct is associated with an 8-bp degenerate barcode sequence (TargetBC). Assuming that the integration sites for each TargetBC are different, we grouped reads from any given replicate that shared the same TargetBC. Based on our observation that array collapse is more frequent than the array expansion, we selected the read with the maximum number of TAPE-monomers from each set of reads that shared a TargetBC. If multiple reads were tied by this criterion, we selected the one (or one of the ones) with the most edits for presentation in **Supplementary Figure 8**. For presentation, we selected reads that have at least 3 insertions and at most 25 TAPE-1 monomers.

