## Supplementary Figures for "A temporally resolved, multiplex molecular recorder based on sequential genome editing"

### Supplementary information for:
### A temporally resolved, multiplex molecular recorder based on sequential genome editing

Junhong Choi[1,2,#], Wei Chen[1,3], Anna Minkina[1], Florence M. Chardon[1], Chase C. Suiter[1,4], Samuel G. Regalado[1], Silvia Domcke[1], Nobuhiko Hamazaki[1,2], Choli Lee[1], Beth Martin[1], Riza M. Daza[1], Jay Shendure[1,2,5,6,#]

[1] Department of Genome Sciences, University of Washington, Seattle, WA 98195, USA
[2] Howard Hughes Medical Institute, Seattle, WA 98195, USA
[3] Molecular Engineering and Sciences Institute, University of Washington, Seattle, WA 98195, USA
[4] Molecular and Cellular Biology Program, University of Washington, Seattle, WA 98195, USA
[5] Brotman Baty Institute for Precision Medicine, Seattle, WA 98195, USA
[6] Allen Discovery Center for Cell Lineage Tracing, Seattle, WA 98195, USA

**Supplementary Figure 1. The relative insertional frequencies of k-mers to DNA Ticker Tape are determined by relative pegRNA abundances as well as by insertion-dependent sequence bias.**

**Supplementary Figure 2. Designing 48 3xTAPE constructs from eight basal CRISPR spacer sequences.**

**Supplementary Figure 3. Testing longer barcode insertions at TAPE-1 and TAPE-27.**

**Supplementary Figure 4. Hierarchical clustering analyses of identified unigram barcodes based on the bigram matrices.**

**Supplementary Figure 5. Barcode distribution in a lineage tracing experiment with 5xTAPE-1.**

**Supplementary Figure 6. Reconstructed lineage trees based on the 5xTAPE-1 insertion patterns.**

**Supplementary Figure 7. Sanger sequencing traces for cloned 12xTAPE-1 and 20xTAPE-1 constructs.**

**Supplementary Figure 8. Recovery of ~12x- and ~20x-TAPE-1 arrays after prolonged editing.**

**Supplementary Table 1. Read counts and editing efficiencies for 2xTAPE-1, 3xTAPE-1, and 5xTAPE-1**. Sequencing reads were grouped based on the observed editing pattern. For example, reads from the 2xTAPE-1 array were categorized into four groups: (a) no edit at either TAPE-1 site (‘OO’); (b) 5-bp insertion at the first TAPE-1 site only (‘XO’); (c) 5-bp insertion at the second TAPE-1 site only (‘OX’); or (d) 5-bp insertions at both TAPE-1 sites (‘XX’). For reads from the 5xTAPE-1 array, editing groups were simplified by further grouping sequential (OOOOO, XO, XXO, XXXO, XXXXO, and XXXXX) and non-sequential (OX, NOX, NNOX, and NNNOX, where N can be either O or X) editing patterns. Editing efficiencies at each site were calculated as the fraction of reads with an edit at the site over the total number of reads in which the site had been activated via insertion of the ‘key’ that completed the spacer sequence. 5-bp insertions were tested except for the 5xTAPE-1 array, where 6-bp insertions (random 3-bp plus 3-bp key sequence) were also tested.

**Supplementary Table 2. Inferred vs. true event orders for Programs 1, 2, and 3.** The order and ratio of events over sixteen epochs (sequential transfections) were inferred using recording to 5xTAPE-1 in a population of HEK293T cells. For each program, the order of events was inferred using unigram information only (‘Unigram order’) or using both unigram and bigram information (‘Bigram order’). In Program-3, two barcoded pegRNAs were co-transfected in each epoch. For Program-1 and Program-2, the correct order across all 16 epochs was identified using bigram information. For Program-3, a single swap between epochs 4 and 5 differentiated the actual order vs. the order inferred using bigram information.

**Supplementary Table 3. Inferred vs. true event orders and frequencies for Programs 4 and 5.** The order and ratio of events over eight epochs (sequential transfections where each transfection introduced a mix of two barcoded pegRNAs) were inferred using recording on 5xTAPE-1. The order of events was inferred using the unigram information alone (‘Unigram order’) or using both unigram and bigram information (‘Bigram order’). For Program-4 and Program-5, the correct order across all 8 epochs was identified using bigram information. Furthermore, the relative ratio at which each pair of barcodes was observed in Program-4 vs. Program-5 agreed well with the relative ratio at which the corresponding pegRNAs were used between the programs.

**Supplementary Table 4. TargetBC-5xTAPE-1 alleles and read counts.** Both 8-bp target barcode (TargetBC) upstream of 5xTAPE1 and 3-bp insertion barcode (InsertBC) with GGA key sequences were extracted from each sequencing reads. Each of the 6644 alleles listed has up to six repeats (due to rare expansion from the expected five repeats) of the TAPE-1 monomer (TGATGGTGAGCACG), followed by an 11-bp sequence (TGATGGTGAGC) that acts as the homology arm for prime editing at the last TAPE-1 target.

**Supplementary Table 5. Shannon entropy and Kullback-Leibler divergence measurements on lineage trees.** For each highlighted TargetBC (GGGGTTTT, ATAGGGCC, AGTTGCGT, GTTTACGC), we calculated the Shannon entropy of InsertBCs for the first three insertion sites, and their Kullback-Leibler divergences (D_KL) against Site-1 insertions across all integrants. For Site-2 and Site-3, we calculated the Kullback-Leibler divergences of each subtree (*i.e.* using read counts from alleles that share InsertBCs at earlier site(s) along the tape) against the Site-1 background distribution, and averaged them across all subtrees. TargetBCs are ordered left-to-right by their Site-1 entropy. stdev stands for calculated standard deviation. For each tree, the number of branches by Site-1, and mean and standard deviation of number of branches by Site-2 and Site-3 subtrees (defined by shared barcodes at earlier site(s) along the tape) are shown.

**Supplementary Table 6. Nucleic acid sequences of experimental constructs.**

**Supplementary Table 7. Primer sequences used in PCR reactions.**

**Supplementary Table 8. TAPE64 encoding of text symbols to 3-mer barcodes**

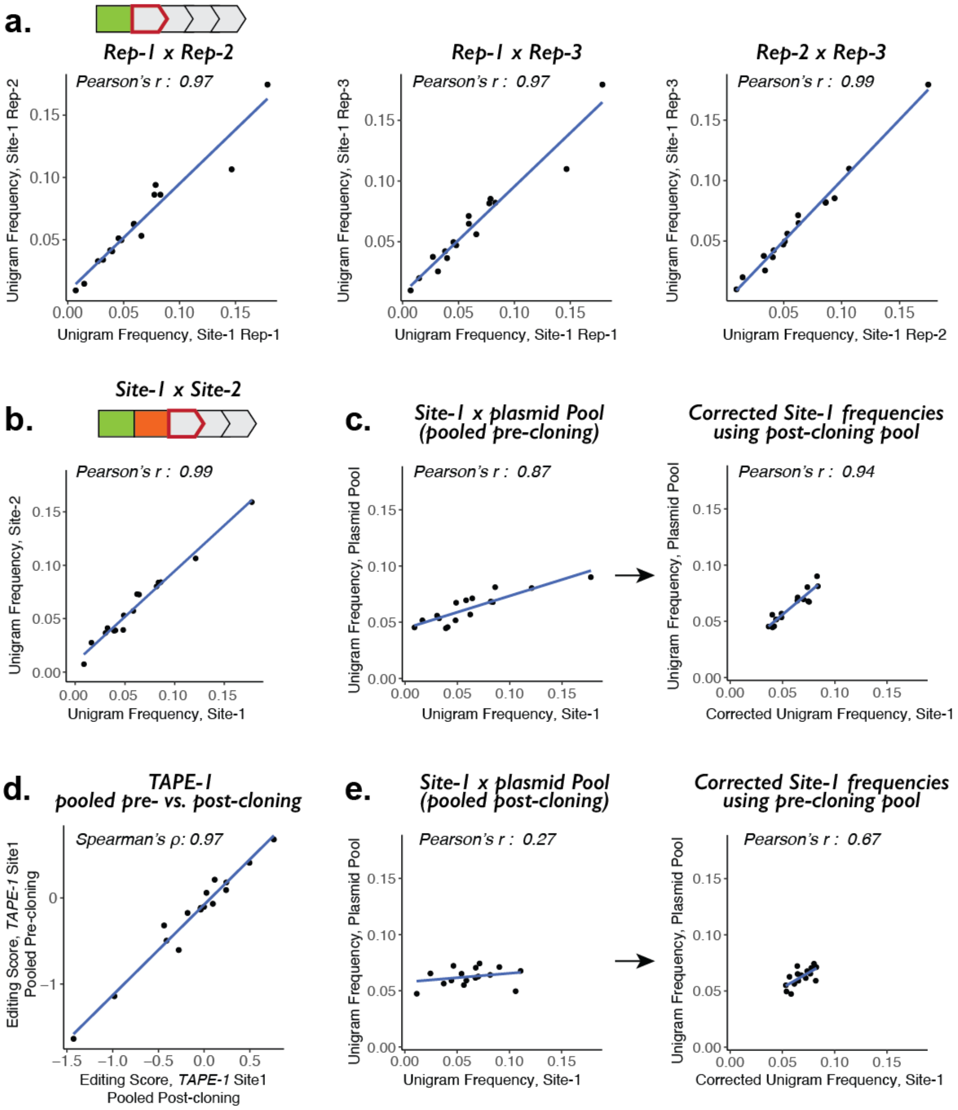


**Supplementary Figure 1. The relative insertional frequencies of k-mers to DNA Ticker Tape are determined by relative pegRNA abundances as well as by insertion-dependent sequence bias. a.** Pairwise scatterplots of unigram frequencies of NNGGA insertions at the initiating monomer of 5xTAPE-1 among three transfection replicates. **b.** Scatterplot of unigram frequencies, averaged across three transfection replicates, at the initiating vs. second monomer of 5xTAPE-1. **c.** Scatterplot of averaged unigram frequencies at the initiating monomer in "pre-cloning pooling" experiment vs. the abundances of NNGGA pegRNA-expressing plasmids (left). Insertional bias was corrected for with data from a separate experiment using NNGGA pegRNA-expressing plasmids that were pooled post-cloning, resulting in a better correlation with the abundances of pegRNAs in the plasmid pool (right). Corrections were done by dividing pre-cloning unigram frequencies by post-cloning unigram frequencies at the initiating monomer and multiplying by post-cloning pegRNA plasmid frequencies. **d.** Scatterplot of NNGGA editing scores calculated on the initiating monomer of the 5xTAPE-1 target edited by pegRNA-expressing plasmids pooled pre-cloning vs. post-cloning. Editing scores for each insertion are calculated as log2 of the ratio between insertion frequencies and the abundances of pegRNAs in the plasmid pool. Spearman's *p* was used instead of Pearson's *r*. **e.** Scatterplot of averaged unigram frequencies at the initiating monomer in "post-cloning pooling" experiment vs. the abundances of NNGGA pegRNA-expressing plasmids (left). Correcting for insertional bias with pre-cloning unigram frequencies improves the correlation (right).

**a.**

```
HEK3   : GGCCCAGACTGAGCACG TGATGGCAGAGGAA   (Base-1)
TAPE-1 : GGATGATGGTGAGCACG TGATGGTGAGCACG   (Design-1)
TAPE-2 : GGATGATGGCAGAGGAG TGATGGCAGAGGAG   (Design-2)
TAPE-3 : GGACCAGGGTGAGCACG CCAGGGTGAGCACG   (Design-3)
TAPE-4 : GACTGATGGTGAGCACG TGATGGTGAGCACG   (Design-4)
TAPE-5 : GATGATGGGTGAGCACC TGATGGGTGAGCACC  (Design-5)
TAPE-6 : GGACTGATGGTGAGACG TGATGGTGAGACG    (Design-6)

FANCF  : GGAATCCCTTCTGCAGC ACCTGGATCGCTTC   (Base-5)
TAPE-25: GAGACCTGGTCTGCAGC ACCTGGTCTGCAGC   (Design-1)
TAPE-26: GAGACCTGGATCGCTTC ACCTGGATCGCTTC   (Design-2)
TAPE-27: GAGATCCGGTCTGCAGC ATCCGGTCTGCAGC   (Design-3)
TAPE-28: GTAACCTGGTCTGCAGC ACCTGGTCTGCAGC   (Design-4)
TAPE-29: GAACCTGGGTCTGCAGC ACCTGGGTCTGCAGC  (Design-5)
TAPE-30: GCAGACCTGGTCTGCAC ACCTGGTCTGCAC    (Design-6)

K14    : GTTGTTGTTCACCTCCT CGATGGCATTCACT   (Base-2)
TAPE-7 : GGACGATGGCACCTCCT CGATGGCACCTCCT   (Design-1)
TAPE-8 : GGACGATGGCATTCACG CGATGGCATTCACG   (Design-2)
TAPE-9 : GGAGTTGGGCACCTCCT GTTGGGCACCTCCT   (Design-3)
TAPE-10: GACCGATGGCACCTCCT CGATGGCACCTCCT   (Design-4)
TAPE-11: GACGATGGTCACCTCCT CGATGGTCACCTCCT  (Design-5)
TAPE-12: GGACCGATGGCACTCCT CGATGGCACTCCT    (Design-6)

PD1    : AACCTCTCGGCTTTCCC GCGCGGCGCCGCCC   (Base-6)
TAPE-31: GGAGCGCGGGCTTTCCC GCGCGGGCTTTCCC   (Design-1)
TAPE-32: GGAGCGCGGCGCCGCCC GCGCGGCGCCGCCC   (Design-2)
TAPE-33: GGACTCTGGGCTTTCCC CTCTGGGCTTTCCC   (Design-3)
TAPE-34: GATGCGCGGGCTTTCCC GCGCGGGCTTTCCC   (Design-4)
TAPE-35: GAGCGCGGAGCTTTCCC GCGCGGAGCTTTCCC  (Design-5)
TAPE-36: GGATGCGCGGGCTTTCC GCGCGGGCTTTCC    (Design-6)

K21    : GGTCCCTCCAGCATCTG CTCTGGCTCCATGG   (Base-3)
TAPE-13: GGACTCTGGAGCATCTG CTCTGGAGCATCTG   (Design-1)
TAPE-14: GGACTCTGGCTCCATGG CTCTGGCTCCATGG   (Design-2)
TAPE-15: GGACCCTGGAGCATCTG CCCTGGAGCATCTG   (Design-3)
TAPE-16: GACCTCTGGAGCATCTG CTCTGGAGCATCTG   (Design-4)
TAPE-17: GTCTCTGGGAGCATCTG CTCTGGGAGCATCTG  (Design-5)
TAPE-18: GGATCTCTGGAGCACCT CTCTGGAGCACCT    (Design-6)

PD2    : GCCTGCAAACTGGTAGG CGCCGGCGTAGGCG   (Base-7)
TAPE-37: GCACGCCGGCTGGTAGG CGCCGGCTGGTAGG   (Design-1)
TAPE-38: GCACGCCGGCGTAGGCG CGCCGGCGTAGGCG   (Design-2)
TAPE-39: GCATGCAGGCTGGTAGG TGCAGGCTGGTAGG   (Design-3)
TAPE-40: GATCGCCGGCTGGTAGG CGCCGGCTGGTAGG   (Design-4)
TAPE-41: GACGCCGGACTGGTAGG CGCCGGACTGGTAGG  (Design-5)
TAPE-42: GCAGCGCCGGCTGTAGG CGCCGGCTGTAGG    (Design-6)

K22    : GCGAGTTCAAGTGCTAC CCGAGGTGCGAGGC   (Base-4)
TAPE-19: GGACCGAGGAGTGCTAC CCGAGGAGTGCTAC   (Design-1)
TAPE-20: GGACCGAGGTGCGAGGC CCGAGGTGCGAGGC   (Design-2)
TAPE-21: GGAAGTTGGAGTGCTAC AGTTGGAGTGCTAC   (Design-3)
TAPE-22: GATCCGAGGAGTGCTAC CCGAGGAGTGCTAC   (Design-4)
TAPE-23: GACCGAGGGAGTGCTAC CCGAGGGAGTGCTAC  (Design-5)
TAPE-24: GGATCCGAGGAGTGCAC CCGAGGAGTGCAC    (Design-6)

PD3    : AGCTGCTCACCACGACG CCAGGGCTGCGGGG   (Base-8)
TAPE-43: GGACCAGGGCCACGACG CCAGGGCCACGACG   (Design-1)
TAPE-44: GGACCAGGGCTGCGGGG CCAGGGCTGCGGGG   (Design-2)
TAPE-45: GGATGCTGGCCACGACG TGCTGGCCACGACG   (Design-3)
TAPE-46: GTCCCAGGGCCACGACG CCAGGGCCACGACG   (Design-4)
TAPE-47: GACCAGGGACCACGACG CCAGGGACCACGACG  (Design-5)
TAPE-48: GGACCCAGGGCACGACG CCAGGGCACGACG    (Design-6)
```

**b.**

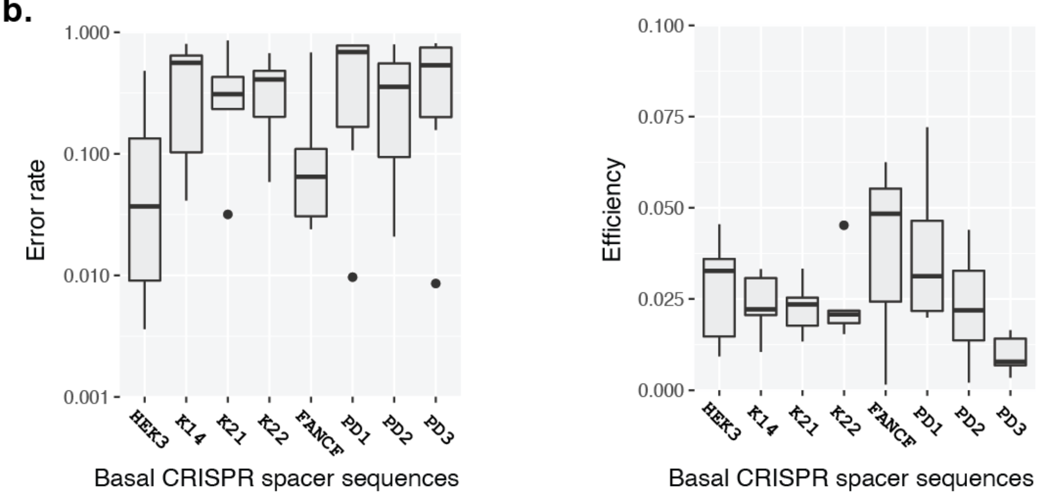


**Supplementary Figure 2. Designing 48 3xTAPE constructs from eight basal CRISPR spacer sequences. a.** For each basal CRISPR spacer sequence (HEK3, K14, K21, K22, FANCF, PD1, PD2 and PD3), six TAPE constructs were derived using six design rules for choosing a 14-bp monomer sequence: Conserving 8 bps towards the 5'-end and 6 bps towards the 3'-end around the Cas9(H840A) nick site (Design-1); Using 13- to 14-bp after the nick site (towards the 3'-end) with G or C at the end (Design-2); Using 14-bp before the nick site, but replacing nucleotide 5 and 6 with 'GG' to generate a PAM sequence (Design-3); Using same 14-bp as Design-1 but altering the 3-bp key sequence (Design-4); Following Design-1 but changing the key sequence length to 2-bp (Design-5) or 4-bp (Design-6), necessitating 15-bp and 13-bp TAPE monomer lengths, respectively. **b.** Boxplots of error rates (left) and efficiencies (right) of 3xTAPE constructs grouped by their basal CRISPR target sequences. Boxplot elements represent: Thick horizontal lines, median; upper and lower box edges, first and third quartiles, respectively; whiskers, 1.5 times the interquartile range; circles, outliers.

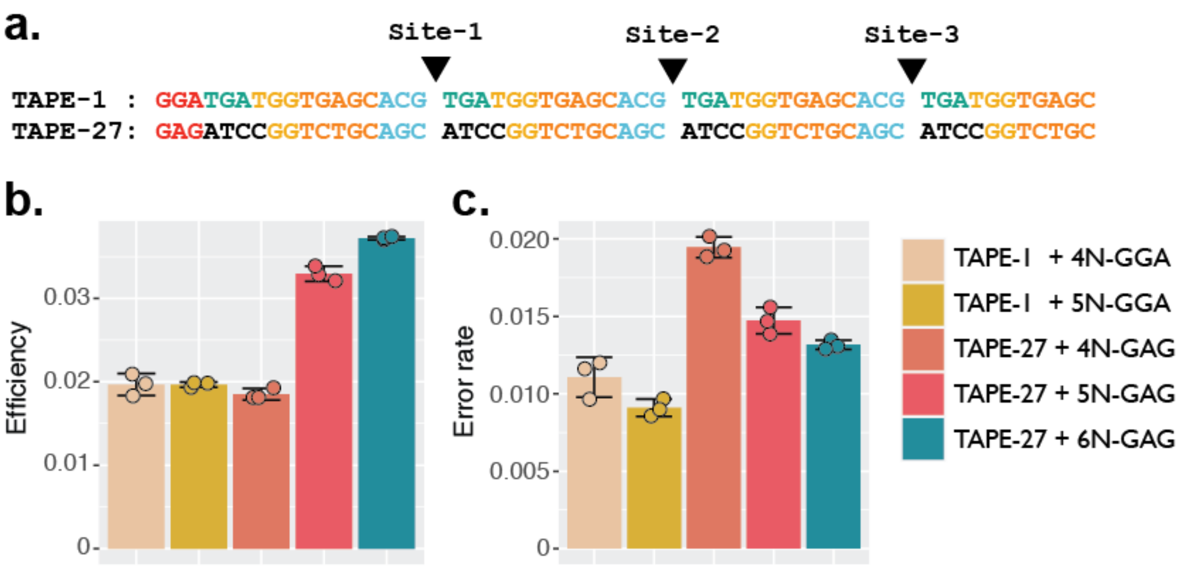


**Supplementary Figure 3. Testing longer barcode insertions at TAPE-1 and TAPE-27. a.** Sequences of 3xTAPE-1 and 3xTAPE-27 arrays. **b-c**. Estimated efficiencies (**b**) and error rates (**c**) measured on TAPE-1 and TAPE-27 for 7-bp, 8-bp, or 9-bp insertions. Insertion sequences include 4- to 6-mer barcodes (synthesized with random N's) and 3-bp key sequences (GGA and GAG for TAPE-1 and TAPE-27, respectively).

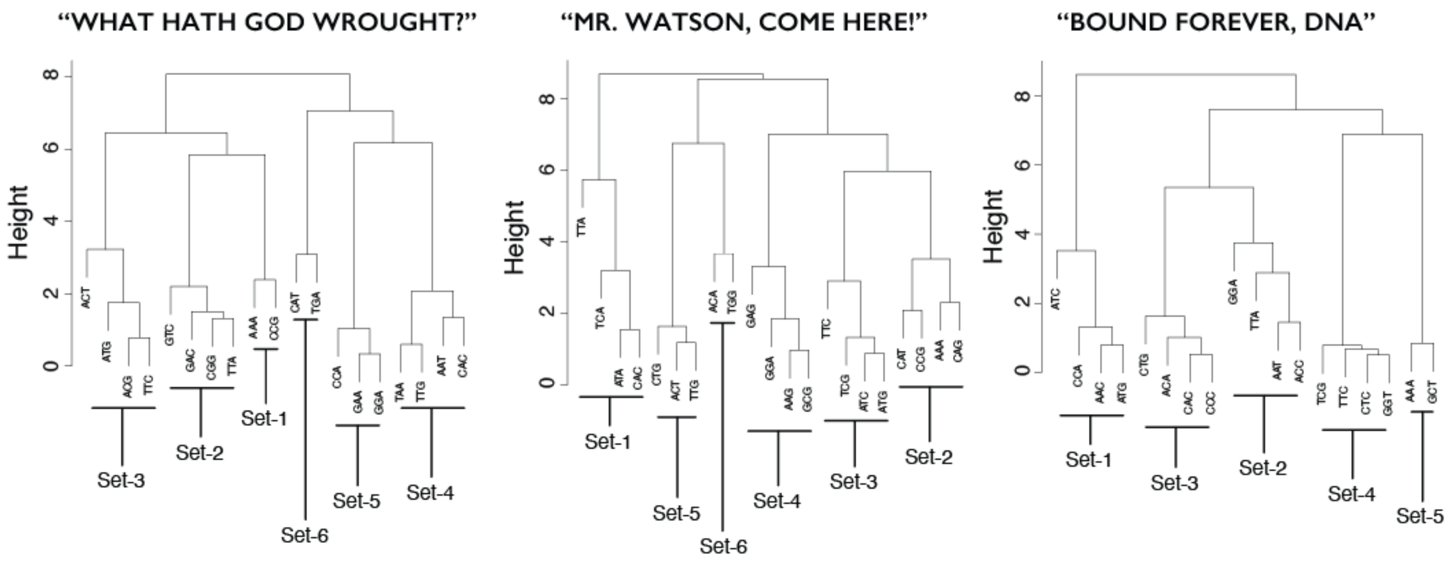


**Supplementary Figure 4. Hierarchical clustering analyses of identified unigram barcodes based on the bigram matrices.** For each message, the normalized bigram matrix was converted to a distance matrix using the euclidean distance measure. The resulting distance matrix was then used for clustering 3-mer barcodes using the complete-linkage clustering method, resulting in a cluster dendrogram for each message. Based on these dendrograms, groups of 2 to 4 barcodes were manually grouped as putative co-transfection sets, and ordered within the set based on unigram frequencies. Sets were ordered relative to one another using the normalized bigram matrix, following the sorting algorithm described in the text and **Figure 4**.

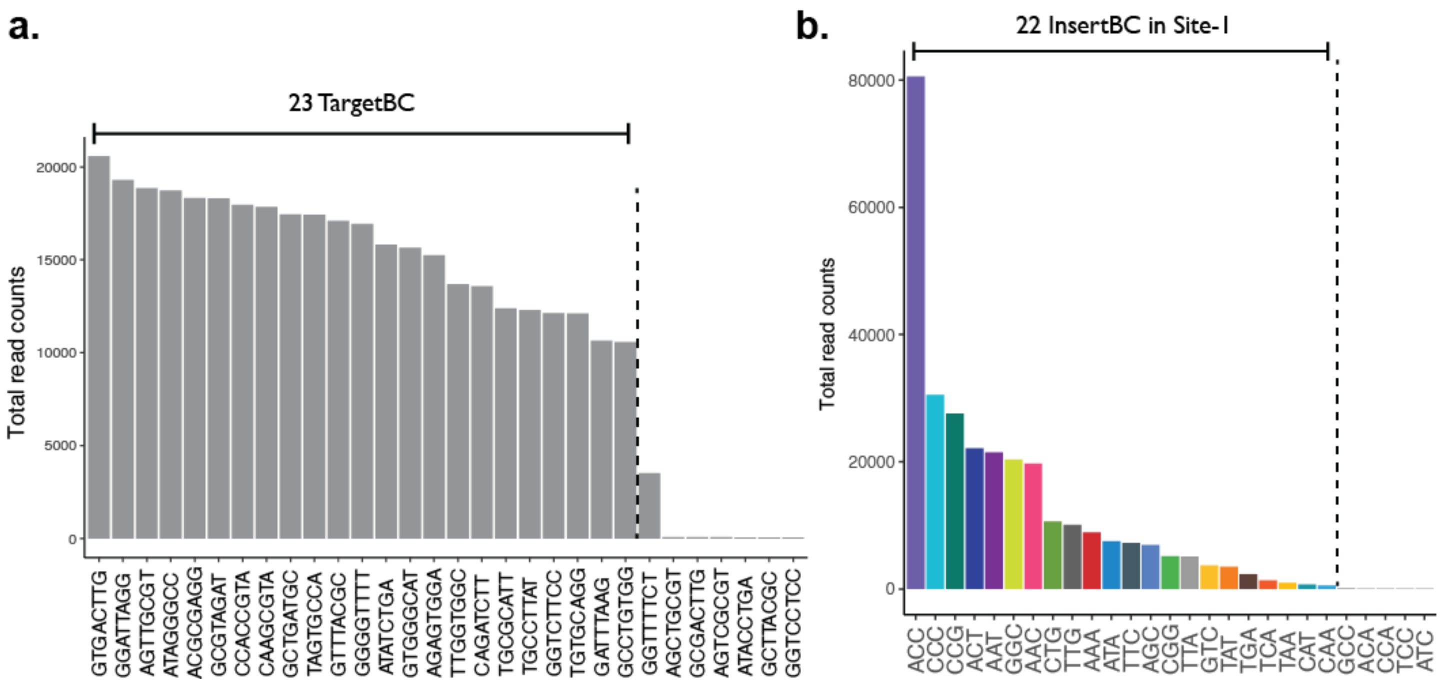


**Supplementary Figure 5. Barcode distribution in a lineage tracing experiment with 5xTAPE-1. a.** Frequencies of TargetBCs observed in all reads. The top 23, or perhaps 24, most frequent TargetBCs are likely to be the actual barcode set in the monoclonal line, whereas less frequent barcodes likely result from sequencing/PCR errors. **b.** Read counts of InsertBC observed in all reads at Site-1 of TAPE-1 arrays. The 22 most frequent InsertBCs are likely to be the actual set encoded by 23 pegRNA integrants in the monoclonal cell line, and are colored the same as tree diagrams shown in **Figure 5c,d** and **Supplementary Figure 6**.

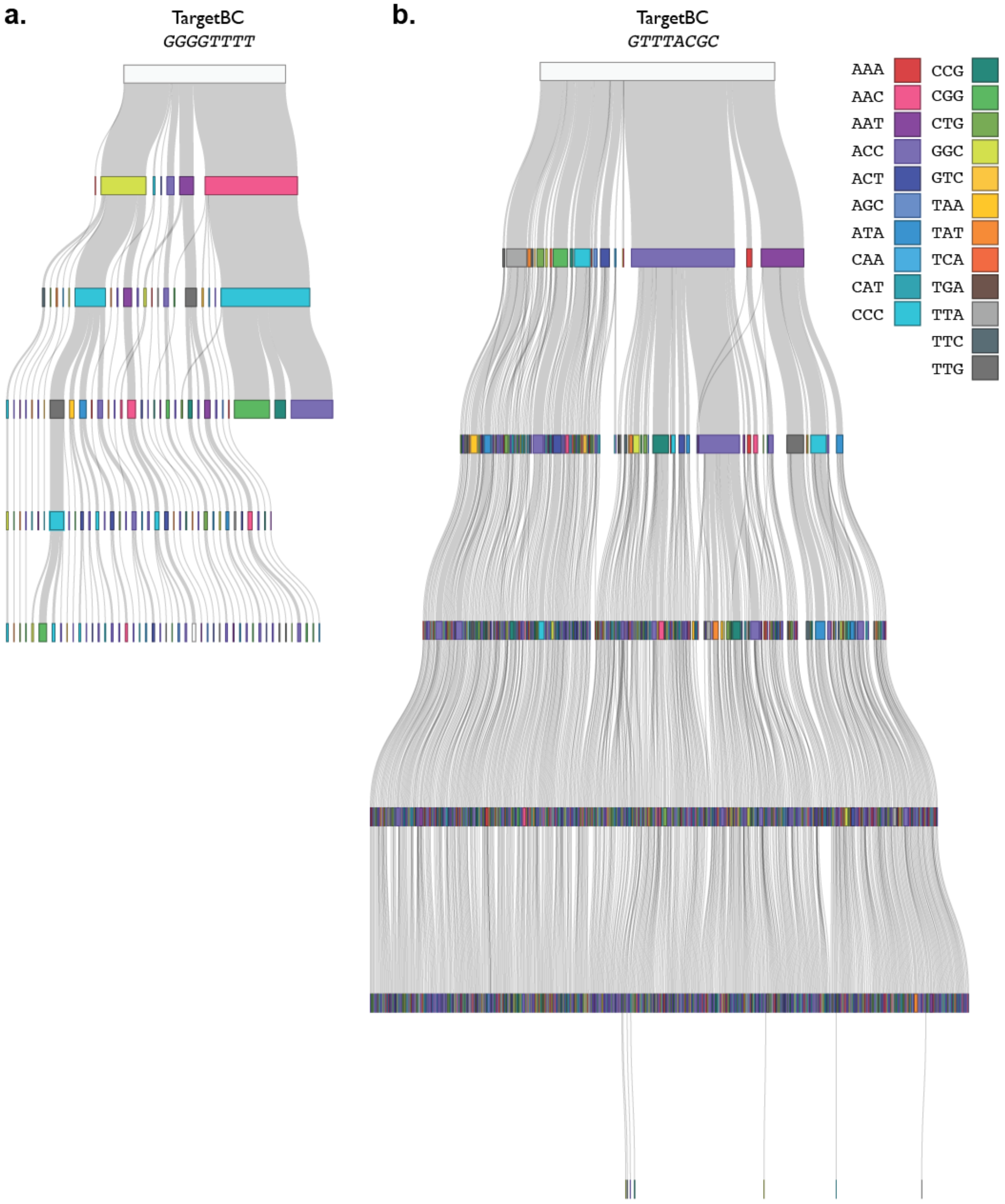


**Supplementary Figure 6. Reconstructed lineage trees based on the 5xTAPE-1 insertion patterns. a.** Reconstruction of editing events for alleles at TargetBC (GGGGTTTT). The TargetBC (GGGGTTTT) reads comprise 66 alleles that range between 3 to 5 repeats of TAPE-1 monomers. The Shannon entropy for the Site-1 insertions is 1.61 bits and $D_{KL}(\textit{Site-1}_{GGGGTTTT}||\textit{Site-1}_{All})$ is 2.18 bits. **b.** Reconstruction of editing events at TargetBC (GTTTACGC). The TargetBC (GTTTACGC) reads comprise 778 alleles that range between 2 to 5 repeats of TAPE-1 monomers. The Shannon entropy for the Site-1 insertions is 2.74 bits and $D_{KL}(\textit{Site-1}_{GTTTTACGC}||\textit{Site-1}_{All})$ is 0.65 bits. The comparatively greater similarity between Site-1 insertions at TargetBC (GTTTACGC) vs. the overall distribution suggests that this tape is being edited more slowly than other tapes, probably due to site-of-integration effects. Consequently, our reconstruction of this lineage tree is more susceptible to errors due to recurrence of the same InsertBC within any given sublineage.

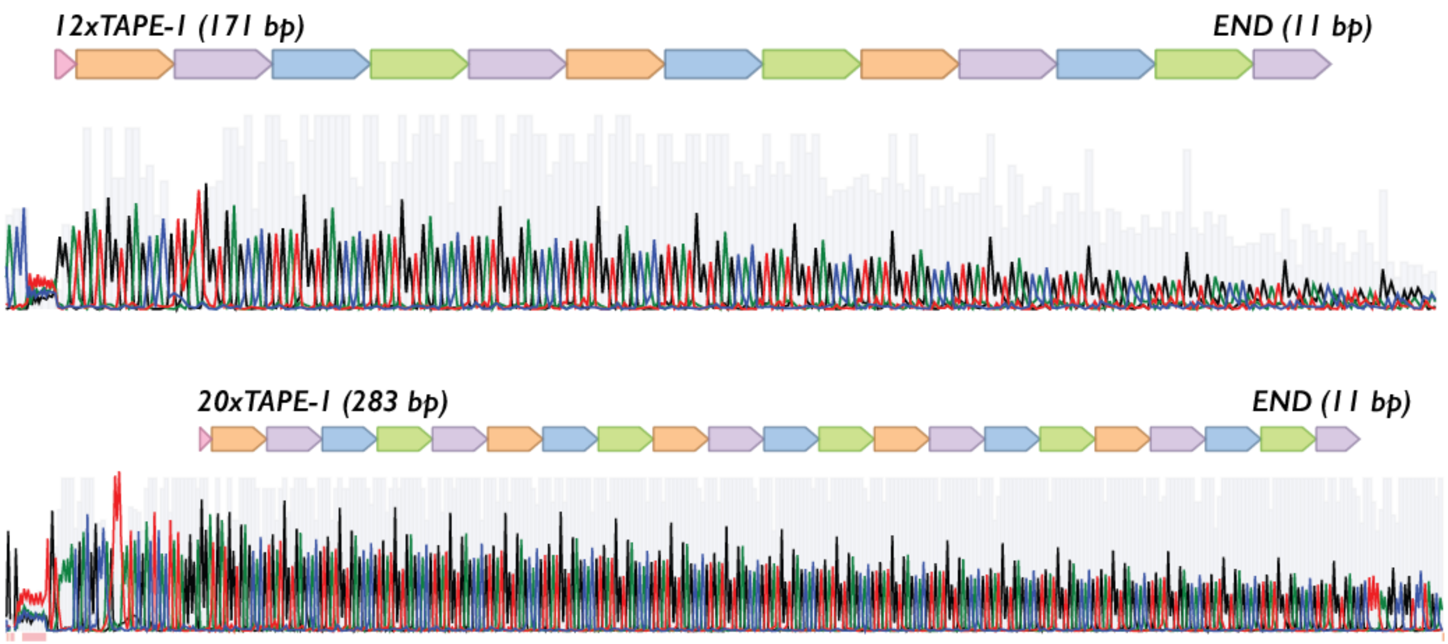


**Supplementary Figure 7. Sanger sequencing traces for cloned 12xTAPE-1 and 20xTAPE-1 constructs.** Each TAPE-array includes 3-bp KEY sequence (GGA for TAPE-1), repeats of 14-bp TAPE-1 monomer, and 11-bp END sequence. Nucleotides A, C, G, and T, in Sanger sequencing traces are colored green, blue, black, and red, respectively. Gray bars in the background are proportional to quality (Phred) for each base call.

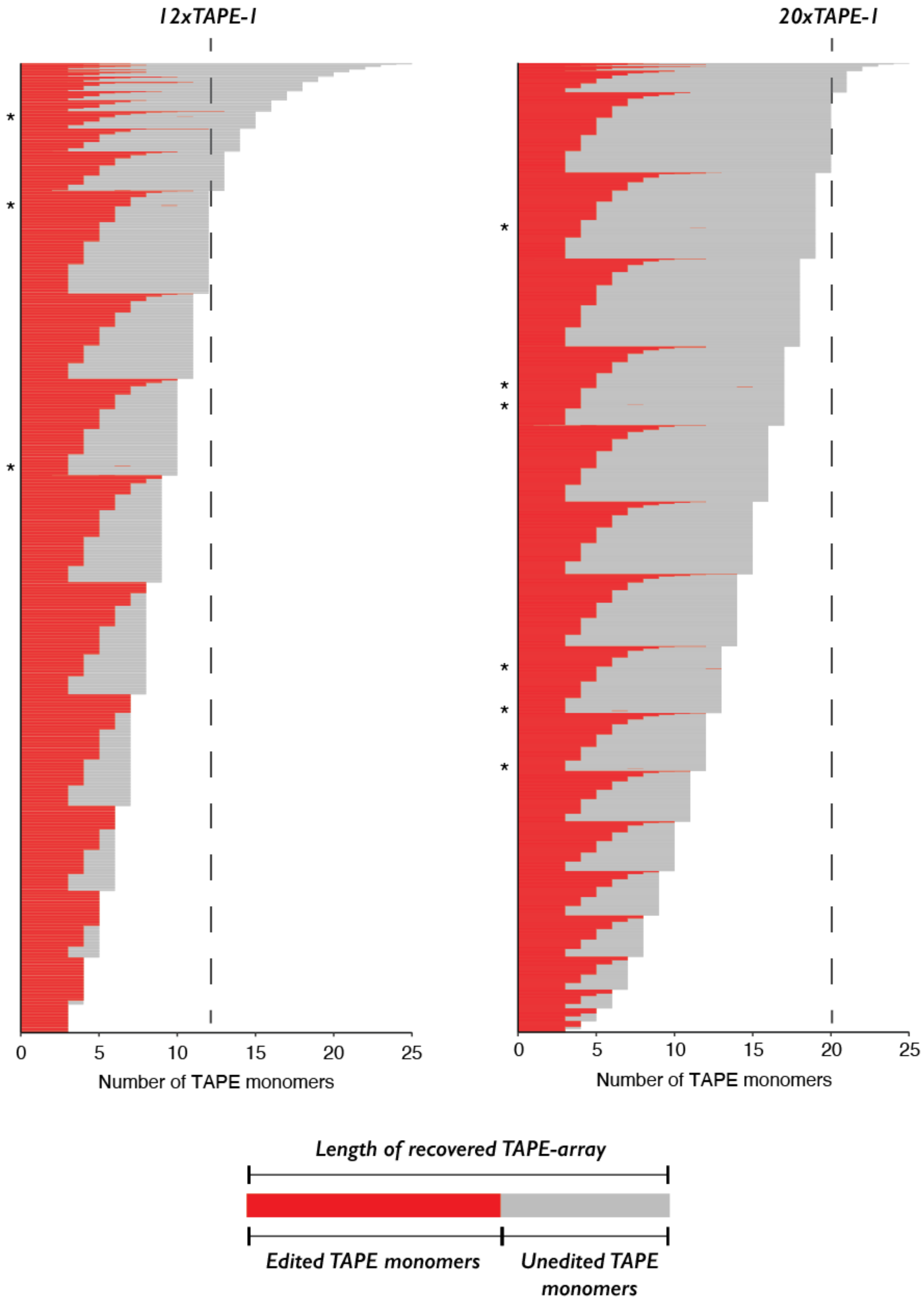


**Supplementary Figure 8. Recovery of ~12x- and ~20x-TAPE-1 arrays after prolonged editing.** ~12x and ~20xTAPE-arrays were integrated into PE2(+) 3N-TAPE-1-pegRNA(+) HEK293T cell line in triplicate, cultured for 40 days for prolonged editing, and recovered via PCR and long-read sequencing on the PacBio platform. Circular consensus sequencing (CCS) reads that had at least 3 NNNGGA insertions and no small indel errors were grouped based on the site of integration (using 8-bp TargetBC barcodes), and a read with the maximum number of TAPE-1 monomers and insertions were selected per TargetBC. Edited portions of each TAPE-array are colored red and illustrate the overwhelmingly sequential, unidirectional editing. Very rarely, we observed non-sequential editing, *e.g.* internal monomers that are edited. These are marked with asterisks to the left of the corresponding row.
